# CMAPS: Causal Mediation Analysis of Perturbation Screens with Application to Genome-scale Perturb-seq Data

**DOI:** 10.64898/2026.05.21.726924

**Authors:** Jingqi Duan, Hyunseung Kang, Sündüz Keleş

## Abstract

CRISPR-Cas9 perturbation screens coupled with single-cell multi-omic profiling enable dissection of gene regulatory mechanisms, yet existing analyses largely quantify total perturbation effects and offer limited insight into the molecular intermediates that transmit these effects. We introduce CMAPS (Causal Mediation Analysis for Perturbation Screens), a semiparametric framework for robust mediation analysis that accommodates unmeasured mediator-outcome confounding and incorporates an adaptive bootstrap test with false discovery rate control. Simulations and data-driven computational experiments show that CMAPS yields accurate, calibrated mediation estimates and robust mediator identification, as confirmed through negative controls and permutation-based validation. Applied to K562 Perturb-seq, CMAPS recapitulates transcriptional cascades downstream of GATA1. In BT16 MultiPerturb-seq data, CMAPS identifies promoter-centric, enhancer-distributed, and mixed *cis*-regulatory programs linking chromatin remodeling factors to transcriptional responses. CMAPS provides a rigorous and interpretable framework for mechanistic inference in single-cell perturbation screens. CMAPS is implemented in R and is available at https://github.com/keleslab/CMAPS.

## Background

The advent of functional genomics using CRISPR-Cas9, combined with single-cell profiling technologies such as scRNA-seq, scATAC-seq, and CITE-seq, has enabled large-scale study of how genetic perturbations impact high-dimensional molecular phenotypes at single-cell resolution, typified by Perturb-seq and its variants. These advancements were impactful in deciphering how genes regulate diverse biological processes and pathway at the single-cell level, uncovering novel gene functions, and revealing the functional consequences of disease-associated perturbations such as cancer mutations and immune regulatory mechanisms [1–9].

With the rapid growth of single-cell CRISPR screens, numerous analytical methods have been proposed (Table A1), each capturing different aspects of perturbation effects but still leaving important gaps. Differential expression analysis [10–16] and distributional comparison approaches [9, 17] identify responsive genes or transcriptome-wide shifts following perturbations. Causal frameworks [18, 19] extend these analyses by estimating direct perturbation effects on individual genes under causal inference assumptions. Moving to a broader scale, gene regulatory network (GRN) inference methods [20–22] aim to reconstruct genome-wide regulator-target maps, providing systems-level views of regulatory architecture. While these approaches hold promise for capturing global causal dependencies, in practice they face challenges such as data sparsity, limited perturbation coverage, and reliance on strong modeling assumptions, which restrict their resolution for pathway-specific mechanisms. Mediation analysis provides a complementary perspective by addressing a more mechanistic question: does a particular molecular intermediate transmit the effect of a perturbation to a downstream gene? Whereas GRNs chart global causal connectivity, mediation explicitly tests pathway-specific hypotheses, offering direct evidence of causal transmission through nominated intermediates.

Mediation analysis decomposes the effect of an exposure *A* on an outcome *Y* into a direct effect and an indirect effect transmitted through a mediator *M*. The classical “product-of-coeffcients” method of Baron and Kenny [23] evaluates these relationships using regression-based tests, while causal mediation analysis formalizes direct and indirect effects within a counterfactual framework [24–28]. When applied to single-cell CRISPR perturbation data such as Perturb-seq [10], causal mediation analysis assesses whether a molecular intermediate *M* (e.g., transcription factors) transmits the effect of a genetic perturbation *A* to downstream phenotypes *Y* (e.g., transcriptomic changes) by decomposing the total perturbation effect. The mediators identified by this approach enable determination of whether a perturbation acts primarily through a specific pathway or via alternative regulatory mechanisms. Accordingly, this framework can be used to address biologically meaningful questions, such as which genes or regulatory elements mediate perturbation effects, which pathways may represent therapeutic targets, and what new regulatory relationships can be hypothesized based on inferred mediation effects and experimentally validated.

Classical mediation analysis assumes no unmeasured confounding of the treatment-outcome, treatment-mediator, and mediator-outcome relationships [23]. Although randomization in single-cell perturbation experiments removes confounding for treatment-mediator and treatment-outcome relationships, it does not eliminate mediator-outcome confounding [28–31]. In single-cell CRISPR screening, such confounding can arise from multiple sources, including variation in sequencing depth, intrinsic cellular heterogeneity [32], cell-cycle state [33], genetic background variation, and transcriptional noise. Some technical factors, such as sequencing depth, can be measured and adjusted for, but many confounders remain unknown or difficult to capture experimentally. Most causal inference methods for single-cell CRISPR data [18, 19] assume no unmeasured confounding, and violation of this assumption can bias effect estimates and inflate false discoveries. To address this limitation, we move beyond the classical mediation framework and adopt semiparametric mediation models [34], which account for unmeasured mediator-outcome confounding.

We developed Causal Mediation Analysis for Perturbation Screens (CMAPS), a statistical framework that enables robust estimation of perturbation effects under unmeasured mediator-outcome confounding, while controlling the false discovery rate and increasing power for mediator discovery (Fig. 1). Applied to genome-scale Perturb-seq in K562 cells [6], CMAPS reveals how perturbing a transcription factor propagates through intermediate TFs to regulate downstream gene expression. For example, upon *GATA1* perturbation, CMAPS recovers TFs with regulatory relationships to *GATA1* (e.g., *STAT5A* and *SP1*) and delineates their regulatory contributions at single-cell resolution. Extending to multiomic settings, application to MultiPerturb-seq in BT16 cells [35] demonstrates how perturbations of chromatin remodelers can influence gene expression indirectly through alterations in *cis*-regulatory element accessibility. This analysis identifies three distinct classes of regulatory programs (promoter-centric, enhancer-distributed, and mixed) and links each class to specific pathway dependencies or potential therapeutic targets.

**Fig. 1:**
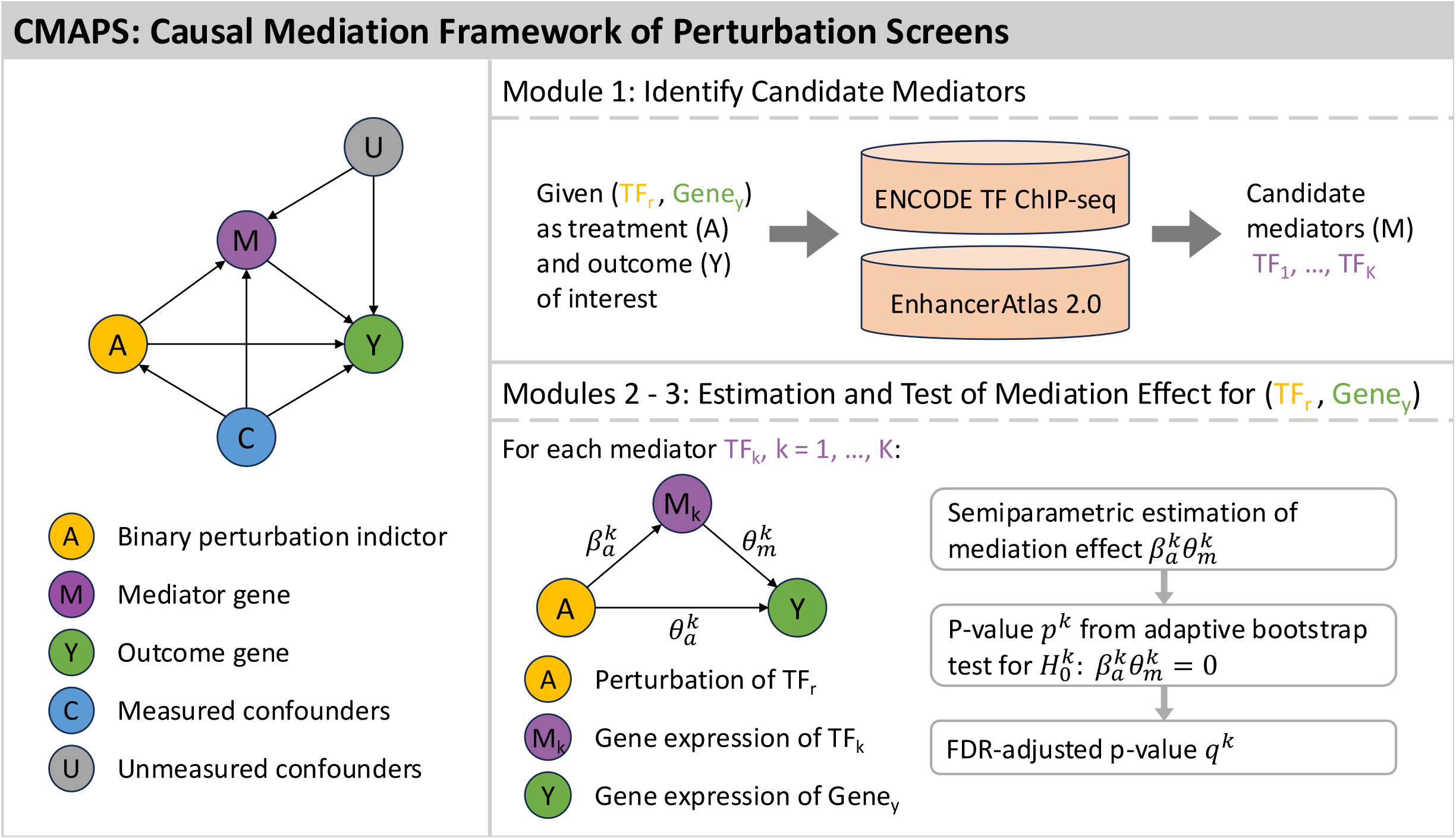
Overview of the CMAPS framework. CMAPS identifies TF-mediator-outcome triplets leveraging public epigenome and enhancer databases from large consortia projects such as ENCODE.

## Results

### Overview of CMAPS

CMAPS is a causal-mediation framework designed for single-cell perturbation experiments (e.g., Perturb-seq [10], CROP-seq [4], SPEAR-ATAC [8], PROD-ATAC [36]). While these screens typically profile many perturbations across cells, CMAPS analyzes each perturbation target separately by restricting to cells receiving that perturbation and contrasting them with non-targeting controls. As shown in Fig. 1, CMAPS evaluates how perturbing a gene *A* affects downstream gene expression (*Y*) indirectly through changes in the expression of intermediate genes (*M*). Here, we illustrate CMAPS using a TF-TF-gene triplet from a Perturb-seq experiment, but the underlying causal-mediation framework is fully general and applies equally to multi-omic single-cell perturbation assays, as described in the final subsection.

Formally, let *A* ∈ {0, 1} indicate the perturbation indicator, with *A* = 1 for cells bearing any sgRNA targeting gene_*r*_ and *A* = 0 for cells with non-targeting control sgRNAs. Let *M, Y* ∈ ℝ denote the molecular phenotype of mediator gene_*m*_ and outcome gene_*y*_, respectively, in gene_*r*_-perturbed and control cells. In this setting, *Y* corresponds to the expression level of gene_*y*_, the outcome of interest, and *M* corresponds to the expression level of gene_*m*_, a candidate mediator for which the causal pathway *A* → *M* → *Y* is hypothesized to hold. Finally, let **C** ∈ ℝ^*p*^ denote measured cell-level confounders such as batch effects, total unique molecular identifier (UMI) counts, mitochondrial gene percentage, and other technical factors, all of which are adjusted for in the analysis.

Given input {*A, M, Y*, **C**}, CMAPS provides unbiased estimates and statistical inference of the mediation effect, the portion of gene_*r*_’s effect on gene_*y*_ that is transmitted through changes in a mediator gene_*m*_ along the pathway *A* → *M* → *Y*. When evaluating multiple candidate mediators (e.g., gene_*m*1_, gene_*m*2_, …) for a given perturbation gene_*r*_ and outcome gene_*y*_, CMAPS provides a mediation effect estimate and *p*-value for each. This enables systematic identification of the specific intermediate genes through which perturbations of gene_*r*_ influence expression of gene_*y*_.

#### Module 1: Identifying candidate mediators based on regulatory evidence

Because causal mediation analysis requires a prespecified pathway *A* → *M* → *Y* (Fig. 1), we restricted both perturbations and mediators to transcription factors (TFs) in our analysis of Perturb-seq data [6]. TFs frequently act in hierarchical regulatory cascades [37–39], making other TFs plausible intermediates between a perturbed TF and a downstream gene. To ensure biological plausibility of the prespecified pathway, we integrated ENCODE TF ChIP-seq data [40, 41] with promoter annotations from Ensembl [42] and enhancer annotations from EnhancerAtlas 2.0 [43] to identify TF-gene regulatory links. For each perturbed TF_*r*_ and outcome gene_*y*_, we defined candidate mediators as TFs (TF_*k*_, *k* = 1,&, *K*) with regulatory evidence for both TF_*r*_ → TF_*k*_ and TF_*k*_ → gene_*y*_ regulation, thereby satisfying the pathway-existence assumption required for causal mediation analysis [24–28].

#### Module 2: Estimating mediation effect under unmeasured mediator-outcome confounding

With a prespecified causal pathway *A* → *M* → *Y* (e.g., TF_*r*_ → TF_*k*_ → gene_*y*_), CMAPS estimates the mediation effect: the component of the perturbation effect of TF_*r*_ on gene_*y*_’s expression that operates through changes in TF_*k*_’s expression. Although perturbations are randomized in single-cell experiments, mediator-outcome confounding may persist, motivating the use of a semiparametric framework that yields valid effect estimation in the presence of unmeasured confounders [34]. In addition to Assumptions 1–3 from classical causal mediation analysis (see Methods), CMAPS requires the following identification conditions [34]:

##### Assumption 4

A4(a) (Semiparametric partially linear model)

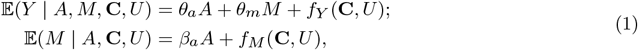

where *f*_*M*_ (*·*) and *f*_*Y*_ (*·*) are unspecified functions of **C** and *U*.

A4(b) (Conditional independence of perturbation) *A* ┴ *U* | **C**.

A4(c) (Heteroscedasticity) v*ar*(*M* | *A* = 1, **C**) ≠ v*ar*(*M* | *A* = 0, **C**).

Here, *U* denotes an unmeasured confounder of the mediator-outcome relationship. A4(a) allows non-parametric dependence of mediator and outcome on measured and unmeasured confounders. A4(b) holds because perturbations are randomly assigned to cells. A4(c) requires that the perturbation change the mediator’s variance. In Perturb-seq, for example, perturbing a TF can alter both the mean and variance of a downstream TF’s expression. Such perturbation-induced changes in the mediator’s distribution provide variation not driven by unmeasured confounders, enabling CMAPS to separate the mediator’s causal effect from bias due to *U*. This assumption can be empirically assessed using heteroscedasticity tests [44].

Under Assumptions 1-4, the natural direct and indirect effects are identifiable [45]:

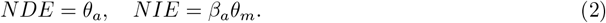

Under standard regularity conditions, the semiparametric estimator is consistent, asymptotically normal (CAN), and multiply robust [34].

#### Module 3: Testing for significant mediators

A key goal of CMAPS is to identify which candidate mediators truly transmit the perturbation effect to downstream genes. External regulatory evidence (Module 1) can nominate many possible mediators for each perturbation and outcome; for example, *GATA1* perturbation in K562 cells yields an average of 66 candidates. Only a subset of these TFs genuinely carries the causal signal, motivating a reliable and powerful mediation test.

For each perturbation TF_*r*_ and outcome gene_*y*_, CMAPS evaluates all candidate mediators TF_*k*_, *k* = 1,&, *K*, individually by testing whether the mediation effect 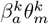 is nonzero:

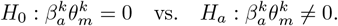

where 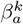 and 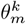 are the treatment-mediator and mediator-outcome effects, respectively, and their product 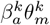 is the mediation effect of TF_*k*_.

Classical tests such as the Sobel test [46] have low power when both coeffcients are close to zero [47–49]. CMAPS addresses this by integrating the semiparametric estimator (Module 2), which remains unbiased under unmeasured mediator-outcome confounding, with the adaptive bootstrap (AB) test [50]. The AB procedure handles the nonstandard behavior of the product-of-coeffcients statistic and provides calibrated p-values across all null settings (see Methods).

#### Extension to multi-omic perturbation screens

Single-cell multi-omic data such as MultiPerturb-seq [35] simultaneously profile chromatin accessibility and transcription, enabling analysis of how perturbations modify chromatin state and, in turn, influence gene expression [51–53]. This motivates the use of mediation analysis to test whether accessibility changes mediate transcriptional responses. To investigate this in a perturbation context, we applied CMAPS to MultiPerturb-seq [35], which jointly measures scATAC-seq and scRNA-seq under CRISPR perturbations of chromatin remodelers. For each perturbation and outcome gene, *M* denotes accessibility at a linked *cis*-regulatory element (e.g., promoter, enhancer, intragenic region) and *Y* denotes its expression level. The pathway *A* → *M* → *Y* captures how perturbation-induced accessibility changes mediate transcriptional responses. By testing mediation across regulatory elements, CMAPS identifies the specific elements through which remodeler perturbations reprogram gene expression.

#### Computational efficiency of CMAPS

CMAPS achieves scalability through two design choices. First, it uses a one-step estimator to obtain the semiparametric estimates without iteratively solving estimating equations, substantially reducing computation while retaining consistency and asymptotic normality. Second, CMAPS replaces the non-parametric bootstrap in the original adaptive bootstrap (AB) procedure with a parametric bootstrap based on the estimator’s asymptotic distribution. This yields nearly identical empirical distributions for the bootstrapped estimates, standardized scores, and AB statistics (Fig. A1a-c), but at a fraction of the computational cost.

These efficiencies make CMAPS practical for large-scale analyses. In the Perturb-seq analysis of *GATA1* perturbation (∼10,000 cells; ∼8,000 outcomes; ∼69 mediators per outcome), CMAPS evaluated over half a million mediator tests. On a 56-core machine with 220 GB RAM, one-step estimation required 25 seconds per outcome on average, and the parametric bootstrap required 4 seconds, demonstrating scalability to high-dimensional mediator sets.

### Simulation benchmarks establish the validity and power of CMAPS

To evaluate the performance of CMAPS, which builds on the proposed semiparametric adaptive bootstrap (`SP-AB`) procedure, we conducted simulations under three generative settings: a single mediator, two mediators, and multiple mediators. We benchmarked `SP-AB` against two existing approaches: (i) the semiparametric estimator [34] with the Sobel test [46] (`SP`), and (ii) the product-of-coeffcients estimator of Baron and Kenny [23] with the Sobel test [46] (`BK`).

Performance was evaluated along three dimensions: (i) bias and precision of mediation effect estimates, (ii) Type I error control under diverse null hypotheses, and (iii) true and false positive rates (TPR and FPR) for identifying true mediators in multiple-mediator settings. The third metrics are especially relevant for Perturb-seq data, where each outcome gene is linked to many candidate mediators (on average 65 per gene, with up to 120 in some cases), most of which have modest mediation effects (approximately 60% fall between -0.1 and 0.1 based on CMAPS analysis of K562 Perturb-seq data). To align with this structure, our simulations varied the number of candidate mediators (30, 50, and 80) and assigned effect sizes of 0.04, 0.1, and 1 to represent small, moderate, and large mediation effects. For completeness, results from the single- and two-mediator settings are reported in Fig. A2 and A4, with additional single-mediator analyses varying sample size and treatment proportion shown in Fig. A3 and A5. We therefore focus on the multiple-mediator setting, which most closely reflects the complexity of Perturb-seq experiments and provides a realistic benchmark for evaluating CMAPS.

#### CMAPS yields unbiased estimates and valid Type I error control

We first evaluated the bias and standard error (SE) of mediation effect estimates across three generative settings. In the single- and two-mediator settings, both `SP-AB` and `BK` produced unbiased estimates when no mediator-outcome confounding was present, though `SP-AB` exhibited slightly larger SEs (Fig. A2 and A3). Under unmeasured confounding, however, `SP-AB` remained unbiased with only modest variance inflation, whereas BK became biased and highly variable. The multiple-mediator setting, designed to reflect the structure of Perturb-seq data, revealed sharper contrasts (Fig. 2a). Here, `SP-AB` maintained near-zero bias in all scenarios (average bias 0.001-0.019). In contrast, `BK` systematically underestimated effects for true mediators and overestimated for null ones, underscoring its inability to separate signal from noise when unmeasured confounding is present.

**Fig. 2:**
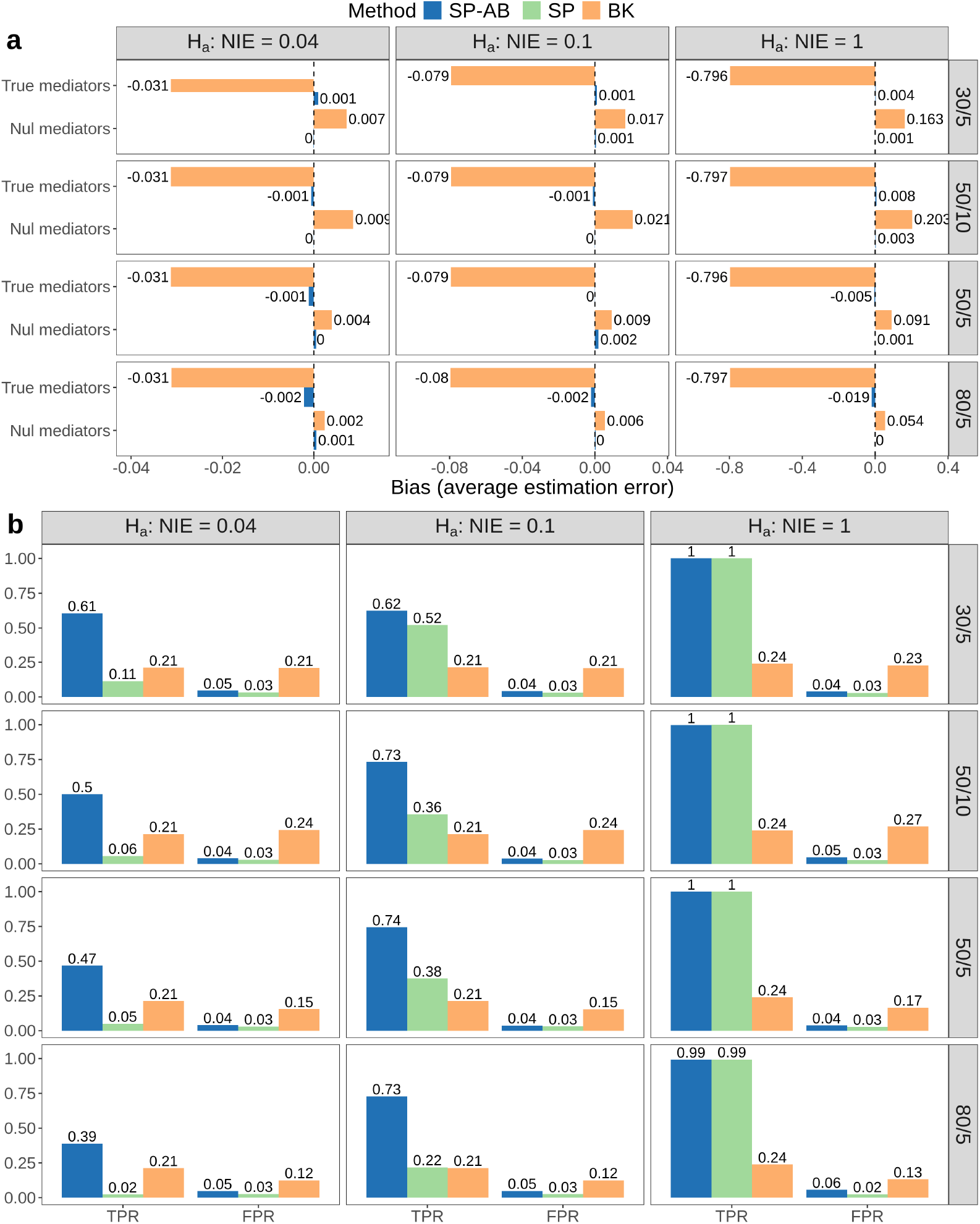
Simulation scenarios in the multiple-mediator setting (Setting 3). Columns indicate the effect size of true mediators, and rows denote the total number of mediators and the number of true mediators (e.g., 30/5 denotes 30 total mediators with 5 true mediators). **a** Average bias of estimated mediation effects, computed separately for true mediators (nonzero effects) and null mediators (zero effects). Bias for each mediator is calculated relative to its true effect and then averaged across mediators within each group. **b** True positive rate (TPR) and false positive rate (FPR) for identifying mediators when both true and null mediators are present. Higher TPR with controlled FPR reflects accurate separation of signal from noise.

We next assessed Type I error control under three null hypotheses: *H*_0,1_: (*β*_*a*_, *θ*_*m*_) = (0, 0), *H*_0,2_: (*β*_*a*_, *θ*_*m*_) = (1, 0), and *H*_0,3_: ((*β*_*a*_, *θ*_*m*_) = (0, 1). In the single- and two-mediator settings, `SP-AB` produced uniformly distributed *p*-values across all nulls, whereas `SP` was conservative under *H*_0,1_ and `BK` was liberal in the presence of confounding (Fig. A4 and A5). Overall, CMAPS delivers unbiased and reliable inference even under unmeasured confounding, making it well suited for Perturb-seq data that feature many candidate mediators with modest effects.

#### CMAPS achieves high sensitivity and specificity for identifying true mediators

In the multiple-mediator setting, data were generated under unmeasured mediator-outcome confounding and included both true mediators, defined as mediators with non-zero mediation effects, and null mediators, characterized by zero mediation effects. A mediator is identified as true if its null hypothesis is rejected at the significance level *α* = 0.05 after multiplicity correction. We quantified performance using the true positive rate (TPR) and false positive rate (FPR) across scenarios that varied both the number of true mediators and their effect sizes (Fig. 2b).

Under this confounding structure, the `BK` method showed poor discrimination, with TPRs and FPRs nearly overlapping (TPRs 0.21 to 0.24; FPRs 0.12 to 0.27). This finding is consistent with the bias patterns observed earlier, where BK underestimated effects for true mediators and overestimated effects for null ones.

We therefore focused on comparing `SP-AB` and `SP`. Both methods consistently controlled Type I error, with FPRs at or below the nominal 0.05 level in all scenarios. As expected, TPRs declined as mediation effects became smaller, reflecting the increased difficulty of detecting weak signals. When effects were large (*NIE* = 1), both methods achieved near perfect detection (TPR close to 1) regardless of the number of true mediators. At a moderate effect size (*NIE* = 0.1), `SP-AB` attained TPRs around 0.7 and clearly exceeded `SP`, whose TPR dropped from 0.52 with 30 mediators to 0.22 with 80 mediators. The contrast was even more pronounced at a small effect size (*NIE* = 0.04): `SP-AB` achieved TPRs of 0.61 and 0.39 with 30 and 80 mediators, respectively, whereas `SP` reached only 0.11 and 0.02 under the same conditions.

These results show that CMAPS, implemented through the `SP-AB` procedure, provides markedly higher sensitivity and specificity for detecting true mediators in settings with many mediators and modest effect sizes, mirroring the challenges of real Perturb-seq data.

### Computational experiments establish the robustness and reliability of CMAPS

#### Empirical validation of true discoveries via permutation tests and negative controls

To assess whether CMAPS captures true mediator signal rather than random variation, we conducted two validation analyses in the *GATA1* Perturb-seq dataset: a negative control analysis and a permutation test. In each case, we computed the discovery rate, the proportion of mediators with FDR-adjusted values below a given threshold *α*. Comparing these rates across the true perturbation, negative controls, and permuted datasets assesses whether observed discoveries exceed those expected under the null. Fig. 3a shows the overall discovery rates at FDR levels *α* = 0.01, 0.02, 0.05, and 0.1. Negative controls and permutations yield discovery rates close to or below the nominal *α*, indicating effective control of false positives. In contrast, the *GATA1* perturbation exhibits substantially elevated discovery rates, particularly at more relaxed thresholds. Fig. 3b displays the distribution of per-outcome discovery rates: null distributions are sharply peaked near zero, whereas the *GATA1* distribution is distinctly right-shifted, indicating many outcomes with substantial mediator signal. Together, these results provide empirical support that CMAPS identifies biologically meaningful mediators rather than artifacts of random variation.

**Fig. 3:**
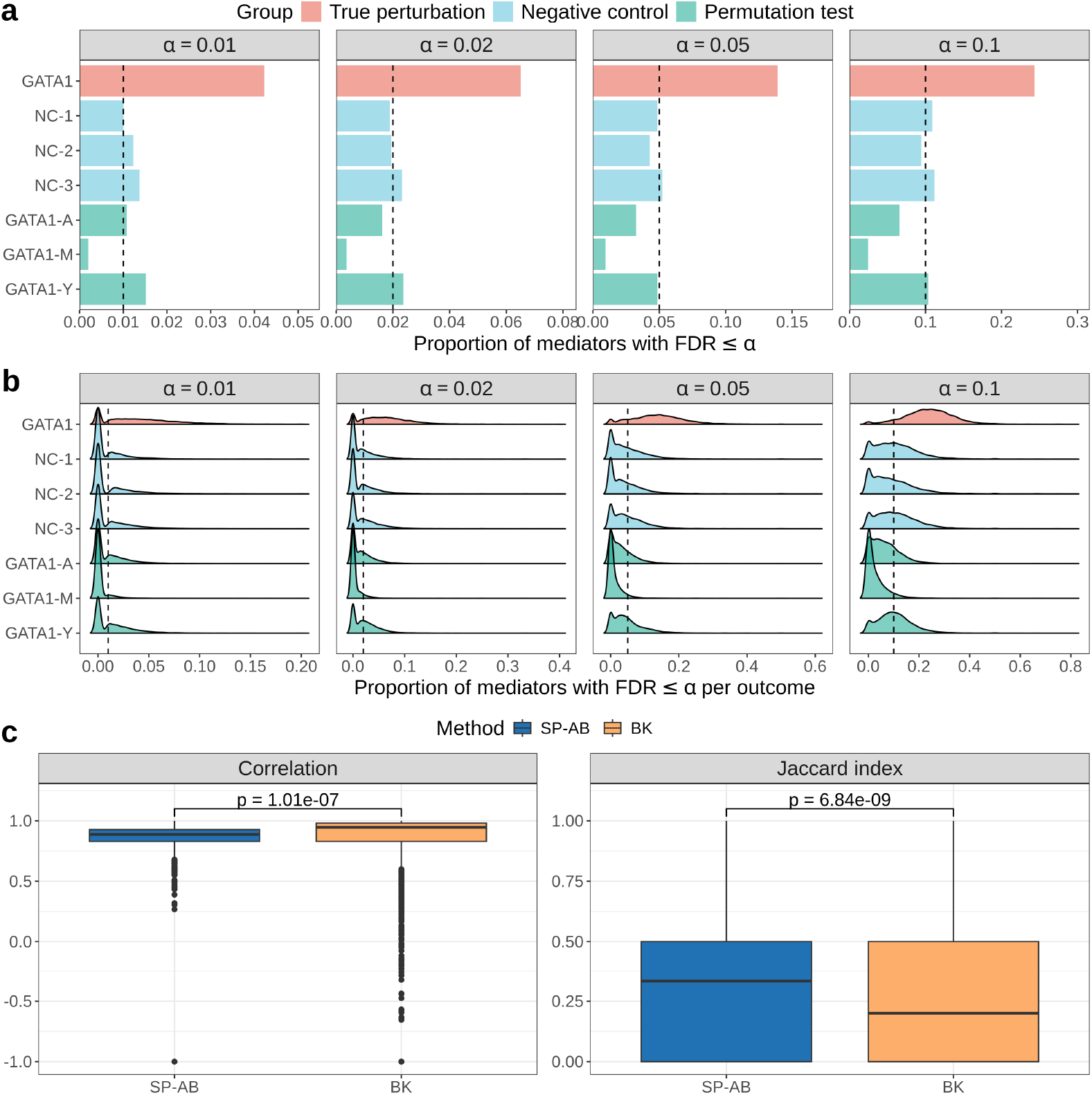
Validation of mediator discovery using negative controls and permutation tests. **a** Discovery rate, defined as the proportion of mediators with FDR-adjusted p-values ≤ α, under three experimental settings: GATA1 perturbation (red), negative controls (blue), and permutation tests (green). Results are shown at four FDR thresholds (α = 0.01, 0.02, 0.05, 0.1; separate column panels). Vertical dashed lines mark the corresponding *ω* level in each panel. **b** Density plots of pergene discovery rates for the same settings and *ω* levels. Distributions for the negative control and permutation analyses are concentrated near zero, whereas the GATA1 distribution is right-shifted, indicating a higher frequency of outcome genes with elevated discovery rates.

#### Robustness of CMAPS to unmeasured mediator-outcome confounding

To assess performance of CMAPS under unmeasured mediator-outcome confounding[54, 55] with a real data computational experiment, we leveraged cell-cycle scores, which summarize each cell’s position in the cell-division program. Cell-cycle scores are generally treated as a potential confounding factor in singlecell analysis because variation in cell-cycle stage drives strong, systematic transcriptional differences that can obscure or falsely inflate biological signals of interest[33].

We treated cell-cycle score as an unmeasured mediator-outcome confounder [54, 55] in the *GATA1* Perturb-seq dataset and compared our proposed `SP-AB` procedure for CMAPS with the classical `BK` approach by fitting both methods with and without adjusting for cell-cycle score. For each outcome gene and its corresponding set of candidate mediators, we quantified agreement between adjusted and unadjusted analyses using two metrics: (i) the correlation between mediation-effect estimates and (ii) the Jaccard index of significant mediators (FDR *<* 0.05). Across outcomes, `SP-AB` achieved substantially higher Jaccard indices than `BK` (Fig. 3c), with differences significant by Wilcoxon, paired-t, and Kolmogorov-Smirnov tests. For correlation, `BK` showed a higher mean value, but exhibited numerous close to zero or negative correlations, whereas `SP-AB` correlations remained uniformly positive (*>* 0.25) and were significantly higher by Wilcoxon and KS tests. These results demonstrate that `SP-AB` is more stable and robust to unmeasured mediator-outcome confounding than the classical `BK` method.

### CMAPS elucidates regulatory mechanisms in K562 cells upon *GATA1* perturbation

Understanding how perturbing a master regulator reshapes downstream transcriptional networks is a core challenge in functional genomics. To investigate this, we applied CMAPS to a genome-scale Perturb-seq dataset in K562 cells, a high-throughput single-cell CRISPR screening platform with scRNA-seq readouts [6]. K562 cells are a human chronic myelogenous leukemia line that retain erythroid features, including expression of embryonic and fetal hemoglobins [56, 57]. These cells rely on the transcription factor *GATA1* to sustain proliferation and erythroid gene programs: loss of *GATA1* disrupts globin and heme-biosynthetic expression and blocks erythroid differentiation [58]. Because *GATA1* regulates cell cycle exit [59], lineage-specific chromatin remodeling [60], and erythroid progenitor survival [61], perturbing *GATA1* provides a powerful test case for dissecting genome-wide regulatory mechanisms.

Within the CMAPS framework, *GATA1* perturbation was evaluated against pooled non-targeting control cells. For each outcome gene, candidate mediators are defined as transcription factors with ENCODE ChIP-seq evidence of binding to that gene [40, 41], yielding an average of 63 mediators per gene (range 1 to 136) across 8,532 outcomes (see Methods). Gene-level expression are used for both the outcome and candidate mediators. This design enables CMAPS to quantify individual mediator’s indirect effects and to pinpoint the key TFs and regulatory mechanisms through which *GATA1* perturbation remodels the transcriptome at single-cell resolution.

#### CMAPS identifies key transcriptional mediators for outcome gene BCL2L1 upon GATA1 perturbation

For a given outcome gene and its candidate mediator set, CMAPS identifies transcription factors whose mediation effects are significant at FDR ≤ 0.05 under *GATA1* perturbation, designating them as key regulators. As an illustrative example, we examined *BCL2L1*, a canonical anti-apoptotic gene, as the outcome gene.

Out of 93 candidate mediators for *BCL2L1*, three TFs showed statistically significant mediation effects (Fig. 4a). The strongest positive mediators, *SP1* and *STAT5A*, bind consensus motifs in the *BCL2L1* promoter [62] and drive its expression under erythropoietin signaling [63]. As a negative mediator, *HMG20B*, a core subunit of the LSD1-CoREST complex, represses anti-apoptotic targets including *BCL2L1* by stabilizing LSD1-GFI1B repressor complexes in erythroid cells [64, 65]. These findings indicate that *BCL2L1* expression is up-regulated through *SP1* and *STAT5A* and down-regulated through *HMG20B* following *GATA1* perturbation.

**Fig. 4:**
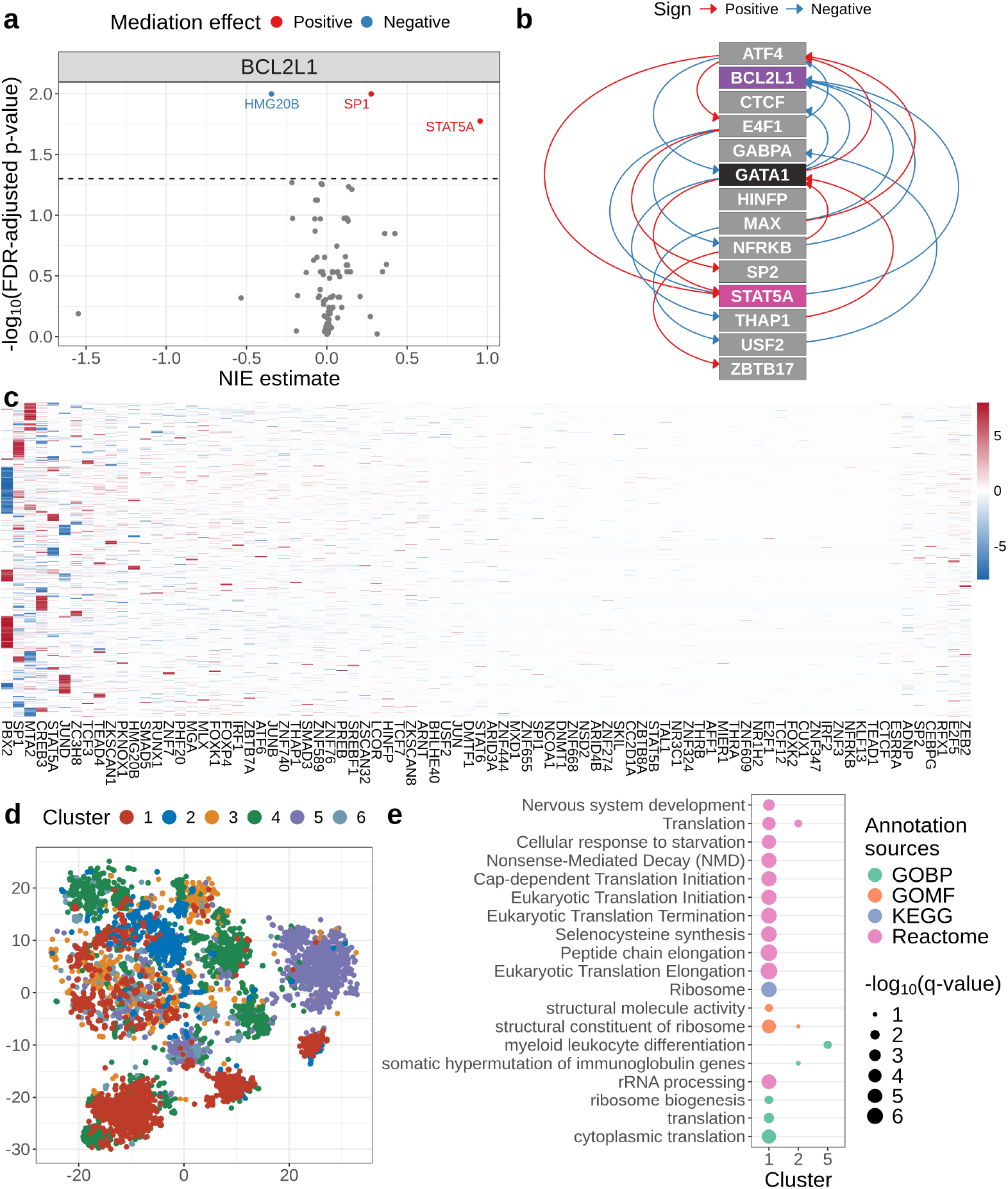
Causal mediation analysis of transcriptional responses to *GATA1* perturbation in K562 cells. **a** CMAPS mediation analysis for *BCL2L1* identifies three significant mediators (FDR ≤ 0.05): positive mediators *SP1* and *STAT5A*, and the negative mediator *HMG20B*; positive effects mean upon *GATA1* loss, *BCL2L1* -expression increases through the mediator. **b** INSPRE-derived causal regulatory relationships among *GATA1*, candidate mediators, and *BCL2L1*; positive values indicate activating relationships and negative values indicate inhibitory ones. **c** Mediation-effect matrix used to cluster genes by shared mediator profiles. **d** Pathway enrichments for each cluster, revealing distinct biological processes engaged after *GATA1* loss. **e** Dot plot of representative enriched annotation terms for each cluster (GO biological process, GO molecular function, KEGG pathway, Reactome pathway). Dot size corresponds to − log 10(q-value) and color indicates annotation resources.

To independently validate these CMAPS results, we applied inspre [21], a causal regulatory network framework, to the Perturb-seq data (see Methods). Inspre recovered a coherent regulatory path *GATA1* → *STAT5A* → *BCL2L1* (Fig. 4b). The direction of regulation along this path agrees with the CMAPS mediation signs, providing orthogonal confirmation of *STAT5A* as a key regulator of *BCL2L1* under erythroid perturbations.

#### Clustering genes by mediator profiles reveals functional modules of GATA1 perturbation

To characterize genes based on their mediator profiles and identify pathways altered by *GATA1* perturbation, we hierarchically clustered the outcome genes (Fig. 4c) and assessed each mediator’s differential impact across clusters. Mediators ranking in the top five for all pairwise cluster comparisons were designated as cluster specific drivers and enrichment analysis was carried out for each cluster.

Cluster 1, driven primarily by *STAT5A* (with *ZNF263* contributing) as a mediator, is strongly enriched in genes for ribosome and translation related processes, including ribosome biogenesis, translation initiation, and peptide chain elongation (Fig. 4d,e). *STAT5A*, the major effector of erythropoietin-JAK2 signaling, promotes transcription of ribosomal protein genes and translation machinery components [66].

Cluster 2, marked by mediators *SP1* and *MTA2*, is enriched in genes for ribosomal structure, core translation functions, and somatic hypermutation of immunoglobulin genes (Fig. 4d,e). *SP1* broadly activates ribosomal protein and elongation factor genes in erythroid cells [67], while *MTA2*, acting through the GATA1-FOG1-NuRD complex, maintains chromatin accessibility at highly transcribed loci [68]. The appearance of somatic hypermutation, normally restricted to AID driven B cell processes, suggests aberrant activation of stress responsive genome editing pathways in K562 cells following *GATA1* perturbation. Together, Cluster 2 mediators appear to coordinate a stress adaptive program linking genome integrity and translational demand.

Cluster 5, with *JUND* as its top mediator, is enriched in genes for myeloid leukocyte differentiation (Fig. 4d,e). *JUND* encodes a JunD subunit of AP-1, a key regulator of myeloid lineage commitment and osteoclastogenesis. AP-1 activity downstream of RANKL-RANK signaling induces NFATc1 and other osteoclast specific genes [69]. Although c-Jun is a dominant driver, *JUND* modulates AP-1 target specificity and contributes to myeloid survival pathways. Its prominence in Cluster 5 suggests that *JUND* promotes the myeloid transcriptional program unmasked by *GATA1* loss.

Clusters 3, 4, 6, and 7 did not show significant pathway enrichment at q *<* 0.05

Overall, these analyses illustrate how causal mediation dissects the diverse regulatory mechanisms downstream of *GATA1* in K562 cells, highlighting both established and unexpected mediators and suggesting future studies on *GATA1*’s partnering transcription factors.

### CMAPS reveals chromatin-mediated regulatory programs in AT/RT cells

We next asked whether CMAPS could extend beyond transcription factor mediators to investigate how genetic perturbations influence gene expression through regulatory elements. Regulatory elements such as promoters and enhancers regulate transcriptional activity, and their function is affected by chromatin accessibility, which is dynamically modulated by chromatin remodelers [70, 71]. To explore this mechanism, we applied CMAPS to MultiPerturb-seq [35], a single-cell multi-omic platform that jointly profiles CRISPR perturbations, transcriptomes (scRNA-seq), and chromatin accessibility (scATAC-seq) within the same cell. Perturbations of chromatin remodelers in this dataset provide a direct way to study mediation along pathways of the form chromatin remodeler perturbation → regulatory element accessibility → gene expression.

MultiPerturb-seq data were generated in BT16 cells, a line derived from atypical teratoid/rhabdoid tumour (AT/RT). AT/RT is an aggressive pediatric brain cancer driven primarily by bi-allelic loss of *SMARCB1*, a core SWI/SNF subunit whose deficiency disrupts chromatin accessibility and transcriptional regulation [72]. The screen profiled 99 epigenetic remodelers across 87,055 cells, including 4,258 non-targeting controls. After quality control (see Methods), 73 perturbations with at least 190 cells each were retained for analysis. Data were obtained from BioProject (accession number PRJNA1160410).

Within the CMAPS framework, each perturbation was evaluated against pooled non-targeting control cells. For each gene, the outcome was its gene-level expression, and candidate mediators were chromatin accessibility measurements at *cis*-regulatory elements (CREs) linked to that same gene. Four CRE classes formed the mediator set: promoter, proximal (downstream, 3’ UTR, and 5’ UTR), distal intergenic, and intragenic (intronic and exonic) (Table 1). This design enabled us to investigate which types of regulatory elements transmit the transcriptional effects of chromatin remodeler perturbations.

**Table 1.**
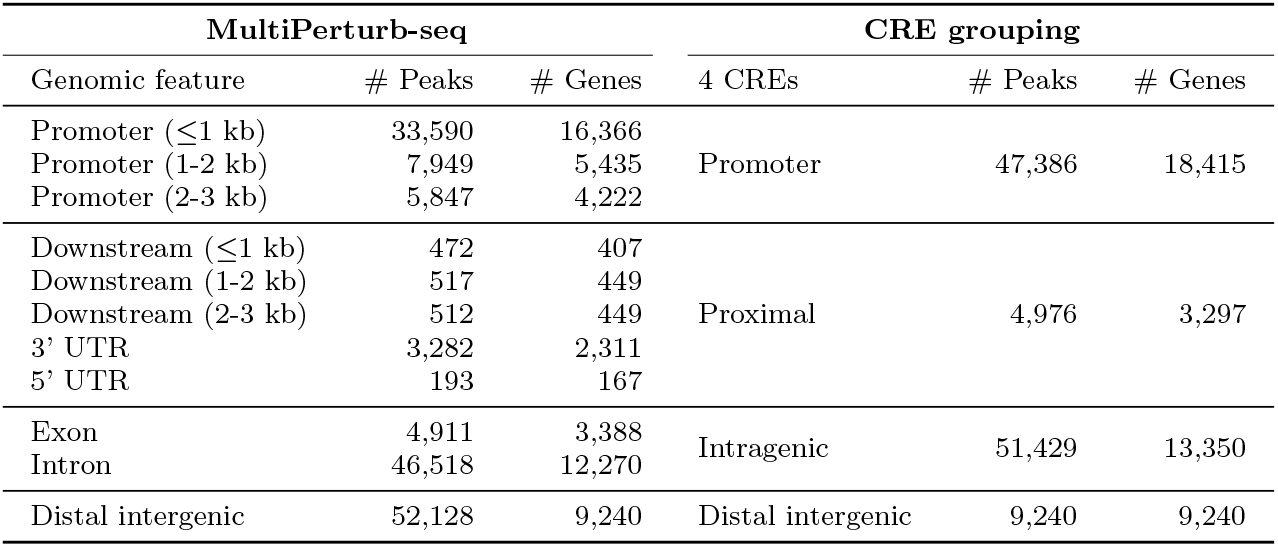
Consolidation of peak-level annotations into four broad cis-regulatory element (CRE) categories. ATAC-seq peaks from the MultiPerturb-seq screen were initially annotated to 11 genomic features and 25,077 genes. For mediation analysis, these features were merged into into four mutually exclusive CRE categories, as indicated. The table lists, for the BT16 AT/RT dataset, (i) the number of peaks assigned to each original genomic feature, (ii) the number of genes harbouring at least one peak in that feature, and (iii) the corresponding peak and gene counts after consolidation into the four CRE categories.

#### Chromatin remodeler perturbations cluster into three mechanistic classes

To investigate whether chromatin remodeler perturbations act preferentially through specific classes of CREs, we quantified the distribution of significant mediation effects (FDR ≤0.05) across promoter, proximal, distal intergenic, and intragenic classes. For each of the 73 perturbations, we computed the proportion of significant CRE-gene pairs assigned to each category, yielding a 73 × 4 proportion matrix with rows summing to one (Fig. 5a).

**Fig. 5:**
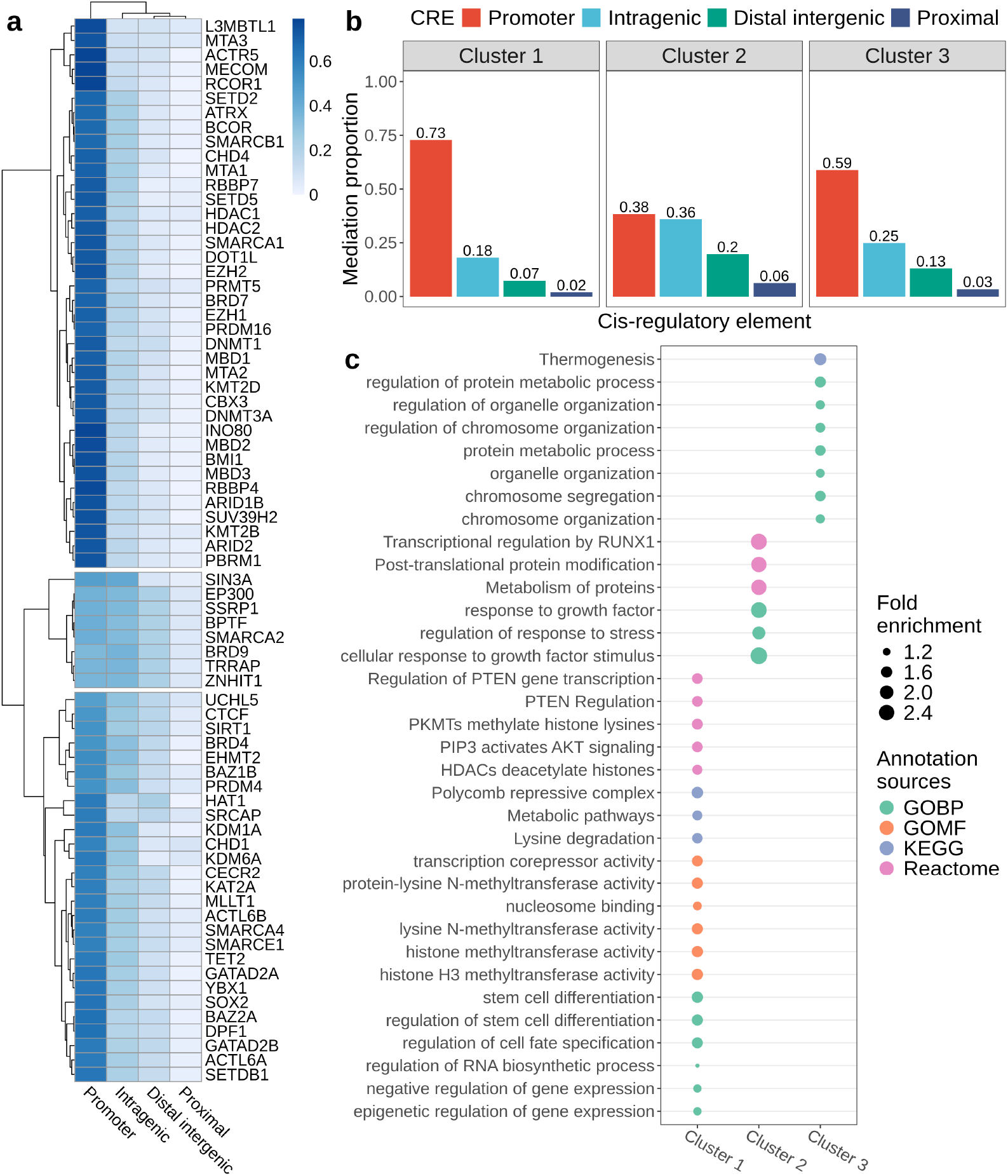
CRE-mediated regulatory landscapes of epigenetic perturbations in AT/RT cells. **a** Heatmap of the proportion of significantly mediated genes (FDR ≤ 0.05) per *cis*-regulatory element (CRE) category (promoter, intragenic, distal intergenic, and proximal) across 73 epigenetic regulator perturbations. Perturbations (rows) and CREs (columns) are clustered by Euclidean distance and complete linkage, revealing three major clusters. **b** Bar plots showing the mean proportion of mediated genes within each CRE category for Clusters 1-3. Cluster 1 is dominated by promoter mediation. Cluster 2 shows balanced promoter and intragenic effects. Cluster 3 exhibits intermediate promoter and intragenic contributions. **c** Dot plot of representative enriched annotation terms for each cluster (GO biological process, GO molecular function, KEGG pathway, Reactome pathway). Dot size corresponds to fold enrichment, and color indicates annotation resources.

Hierarchical clustering of this proportion matrix revealed three groups of chromatin remodelers with distinct CRE mediation profiles, and silhouette analysis supported three clusters as the optimal partition (average silhouette width = 0.51). The distribution of CRE-specific mediation differed strongly across clusters (*χ*^2^ = 2119.1, df = 6, *p<* 2.2 × 10^−16^), demonstrating that remodelers use different regulatory routes to transmit their transcriptional effects. Binary heatmaps of CRE peak presence (Fig. A6d) showed no systematic variation across perturbations, confirming that these profiles reflect biological mechanisms rather than peak availability.

The three clusters, shown top to bottom in Fig. 5a and summarized in Fig. 5b, display distinct mean CRE mediation signatures. Cluster 1 (38 perturbations; 73% promoter, 18% intragenic) is promoter-dominated, suggesting regulation concentrated at core promoters with limited enhancer involvement. Cluster 2 (8 perturbations; 38% promoter, 36% intragenic, 20% distal intergenic) exhibits the strongest enhancer contribution. The comparable promoter and intragenic contributions, together with a distal component, indicate regulation dispersed across gene-internal and distal enhancers rather than confined to promoters. Cluster 3 (27 perturbations; 59% promoter, 25% intragenic, 13% distal intergenic) shows an intermediate pattern that integrates promoter-proximal and enhancer-mediated regulation. Together, these CRE mediation signatures delineate distinct routes through which chromatin remodelers reconfigure transcription in AT/RT cells and motivate the subsequent enrichment analyses.

#### Functional enrichment reinforces cluster-specific regulatory programs

To determine whether the three mechanistic classes correspond to distinct biological functions, we performed enrichment analysis for each CRE-mediation cluster using the 73 perturbed genes as the background across GOBP, GOMF, KEGG, and Reactome pathways (Fig. 5d; see Methods). We also assessed differences in therapeutic relevance.

Cluster 1 is promoter-dominated and enriched for Polycomb activity, histone methyltransferases, and transcriptional and stem cell programs. It is also enriched for PTEN-AKT signaling, which supports survival and growth of rhabdoid tumor cells [73]. Notably, this cluster contains the most clinically advanced targets, such as *EZH2* [74–77], *DNMT1* [78–81], and *HDAC1/2* [82–85], all of which have inhibitors approved for other cancers and are now being evaluated in pediatric brain tumors including AT/RT [86–93].

Cluster 2 shows the strongest enhancer involvement. Enrichment analysis identified growth-factor and stress-response pathways, consistent with studies showing that enhancers involved in stimulus-induced transcription programs [94, 95]. This cluster includes remodelers with emerging therapeutic potential for AT/RT: targeting *BRD9* [96–98] or *EP300* [99] suppresses *SMARCB1* -deficient tumor growth and has progressed to early-stage clinical evaluation [100, 101]. The original MultiPerturb-seq study identified *ZNHIT1* as a candidate target, as its loss was shown to drive chromatin reorganization and differentiation in AT/RT cells [35].

Cluster 3 is enriched for chromosome organization, segregation, and organelle organization, pointing to regulators of genome architecture and chromosomal stability. This broad functional signature is consistent with its mixed CRE mediation profile, in which chromatin remodelers act through diverse regulatory routes rather than a single dominant mechanism. Unlike Clusters 1 and 2, Cluster 3 lacks established therapeutic targets, with evidence limited to preclinical studies showing that inhibition of *BRD4* [102], *EHMT2* [103], and *KDM1A* [104] can suppress tumor growth.

In summary, the three clusters represent distinct mechanistic classes with different biological and therapeutic profiles. Cluster 1 is promoter-dominated and contains remodelers that already serve as therapeutic targets, whereas Clusters 2 and 3 capture enhancer-associated and chromatin-organizing remodelers with emerging or preclinical potential. The clustering framework also highlights additional remodelers with similar regulatory features that may warrant exploration as future therapeutic candidates in AT/RT.

## Discussion

We proposed a causal mediation framework to study the mediating pathways of perturbed genes from a perturbation experiment. Specifically, the semiparametric estimator quantifies both the direct and mediation effects of transcription factors in the presence of unmeasured mediator-outcome confounding, while the adaptive bootstrap test controls type I error and demonstrates high statistical power. Our CMAPS analysis in *GATA1* -perturbed K562 cells yields both gene-specific and global insights into how transcriptional networks are modulated.

For the anti-apoptotic gene *BCL2L1*, we identified three key mediators: *SP1,STAT5A*, and *HMG20B*, Notably, a causal regulatory network inferred from the same dataset independently recovered the pathway *GATA1* → *STAT5A* → *BCL2L1*, demonstrating CMAPS’ ability to pinpoint both known regulators and their mechanistic ordering. When extended genome-wide, CMAPS ranked *SP1, STAT5A, MTA2, PBX2*, and *TEAD4* as the top TFs by the number of genes being mediated, reaffirming their central roles in erythroid differentiation and homeostasis, while also highlighting stress-responsive and RNA-regulatory factors as emerging contributors. Finally, hierarchical clustering of mediation profiles partitioned the transcriptome into distinct functional modules, including *JUND* -driven myeloid and osteoclast programs, *SP1* /*MTA2* -led translation and hypermutation signatures, and *STAT5A*-dominated ribosome biogenesis with novel involvement of selenocysteine synthesis and nonsense-mediated decay pathways. Together, these results illuminate discrete co-regulatory circuits underlying *GATA1*’s genome-wide regulatory impact.

Our *cis*-regulatory element (CRE)–resolved analysis, applying CMAPS to MultiPerturb-seq data, reveals that chromatin remodelers in AT/RT cells segregate into three mechanistic classes defined by how they route transcriptional effects through the regulatory genome. These remodelers differ markedly in the relative contributions of promoter, intragenic, and distal intergenic elements, indicating that the observed profiles reflect intrinsic regulatory strategies rather than differences in chromatin coverage. The promoter-dominated class is enriched for Polycomb and histone-modifying factors with established roles in transcriptional repression and therapeutic relevance, whereas the enhancer-enriched and mixed classes comprise regulators associated with stress-responsive programs, genome organization, and chromosomal stability. Notably, these CRE mediation signatures align with pathway-level enrichment and therapeutic maturity, suggesting that the manner in which a remodeler engages the regulatory genome is closely linked to its biological function and translational potential. Together, these findings support a model in which chromatin remodelers exert their influence through distinct regulatory routes, providing an organizing principle for interpreting perturbation effects and prioritizing candidates for further functional and therapeutic investigation in AT/RT.

One limitation of the proposed framework is that it does not explicitly model multiple mediators simultaneously. In the current implementation, treatment-outcome models are fitted with one mediator at a time, followed by correction for multiple testing across mediators. Although we have extended the semiparametric estimator of [34] to accommodate multiple mediators, this extension remains relatively conservative in terms of statistical power. In particular, while the adaptive bootstrap test effectively controls the Type I error rate, it is inherently designed to evaluate one mediator at a time and does not fully leverage potential joint or synergistic effects. More broadly, biological regulatory systems often involve coordinated activity among multiple mediators within shared pathways or complexes, and analyzing mediators in isolation may obscure higher-order interactions and distributed mechanisms of regulation. An important direction for future work is therefore to develop mediation approaches that can jointly estimate and test multiple mediators, enabling a more comprehensive and mechanistically faithful characterization of complex regulatory networks.

## Methods

### CMAPS algorithm

Algorithm 1 summarizes the semiparametric-adaptive bootstrap (`SP-AB`) procedure.

#### Algorithm 1 Semiparametric-adaptive bootstrap algorithm

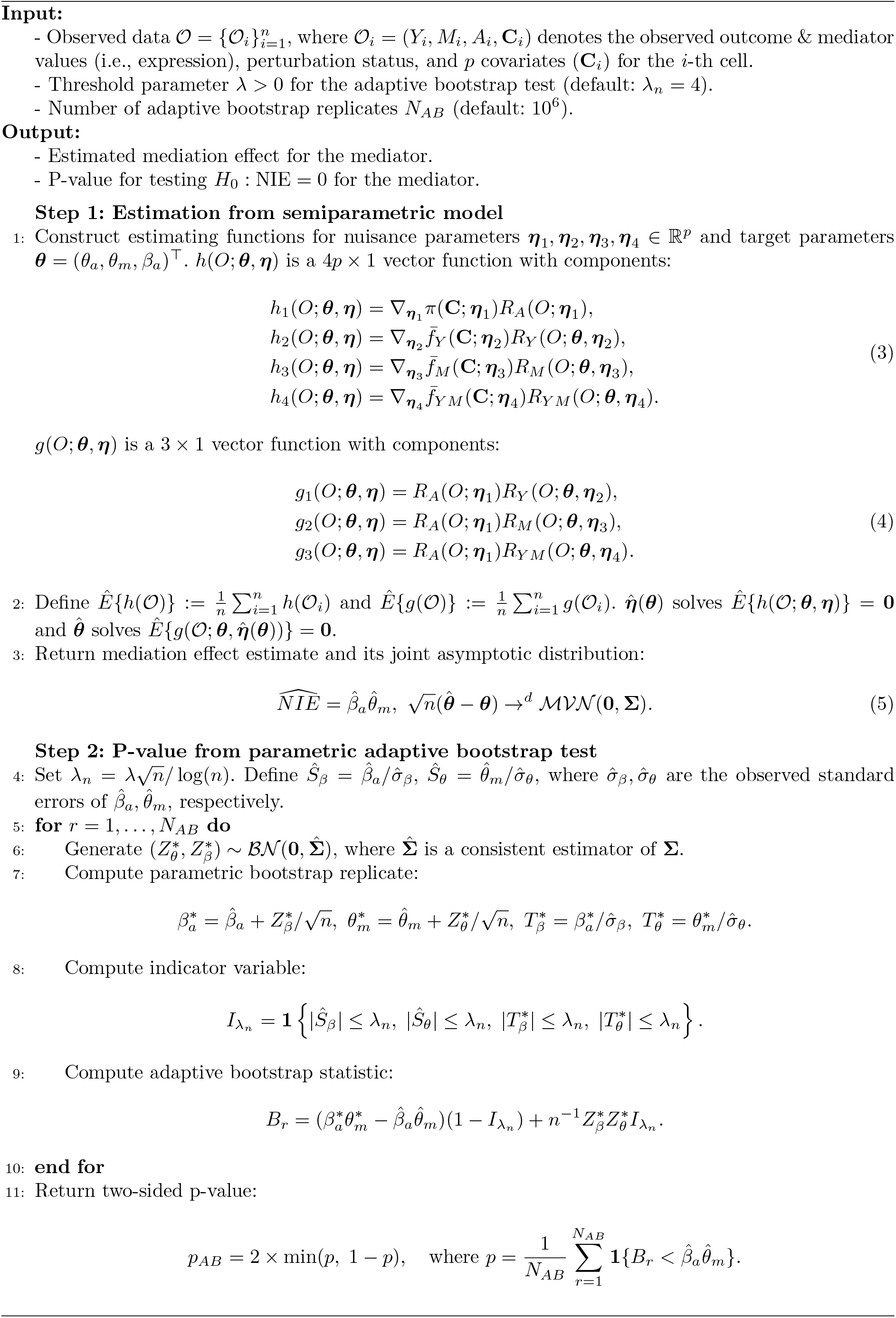

Following functions are used in estimating functions (3) and (4). Note that 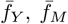, and 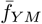 are defined in (9) and (10).

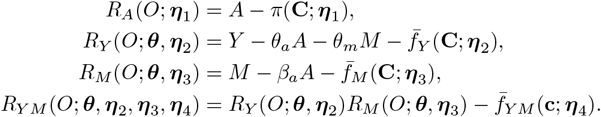

According to Lemma 2 [34], under regularity conditions, 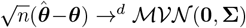as *n* → ∞, where

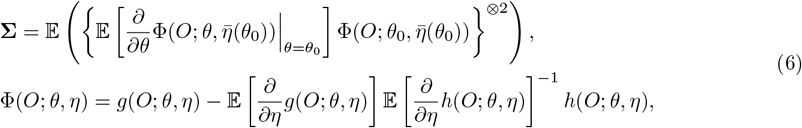

and 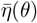 denotes the probability limit of 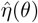. A consistent estimator of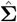 of 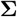 can be constructed by replacing all expected values with empirical mean evaluated at 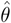 and 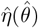.

### Definitions and notations for causal mediation analysis

Let *M* (*a*) denote the counterfactual value of the mediator when the perturbation status is set to *a*, and let *Y* (*a, m*) denote the counterfactual outcome when the perturbation status is set to *a* and the mediator is set to value *m*. More generally, *Y* (*a, M* (*a*^*\**^)) represents the counterfactual value of the outcome when the perturbation status is set to *a*, and the mediator is set to the value it would take under perturbation status *a*^*\**^. For example, using scPerturb-seq data, let *M* and *Y* represent the gene expression of *KLF1* and *HBA1*, respectively, in control versus *GATA1* -perturbed cells. Then, *M* (1) corresponds to the expression of *KLF1* if, contrary to fact, *GATA1* were perturbed. Likewise, *Y* (1,*M* (0)) corresponds to the expression of *HBA1* if *GATA1* were perturbed, but *KLF1* had the expression it would exhibit under control conditions.

With the counterfactual outcomes, we define the following effects of perturbation: the total effect (i.e., TE), the natural direct effect (i.e., NDE), and the natural indirect effect or mediation effect (i.e., NIE).

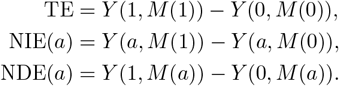

In the context of the *GATA1* perturbation described above, the total effect represents the overall impact of perturbing *GATA1* on the expression of *HBA1*. The natural direct effect (NDE) captures the effect of *GATA1* perturbation on *HBA1* expression when the expression of *KLF1* is held fixed at its counterfactual value. In contrast, the natural indirect effect (NIE) quantifies the effect of changes in the counterfactual expression of *KLF1* on *HBA1*, while holding the perturbation status constant. A positive NIE indicates that perturbing *GATA1* increases *HBA1* expression through its effect on *KLF1*, whereas a negative NIE indicates a decrease in *HBA1* expression mediated through *KLF1*. The total effect can be decomposed into the natural direct and indirect effects as TE = NIE(1) + NDE(0) [27, 28].

### Identification of candidate mediators

Because causal mediation analysis defines the indirect effect relative to a prespecified causal pathway *A* → *M → Y* in the assumed causal diagram [27, 28], where the perturbation *A* influences the mediator *M*, which in turn affects the outcome *Y*, we restricted both perturbations and mediators to transcription factors (TFs) in our analysis of Perturb-seq data [6]. TFs act as master regulators in hierarchical cascades: a primary TF activates secondary TFs, which subsequently regulate downstream genes [37–39]. To ensure that the mediator lies on a plausible biological path between the perturbed TF and the outcome gene, we required ChIP-seq-based binding evidence consistent with a TF_*r*_ → TF_*m*_ → gene_*y*_ relationship. Specifically, for each perturbed TF_*r*_ and outcome gene_*y*_, bound by TF_*r*_, we defined candidate mediators TF_*m*_ as TFs *bound by* TF_*r*_ and *binding* gene_*y*_. Under such architectures, perturbation of TF_*r*_ is expected to alter the expression of TF_*m*_, which modulates the expression of gene_*y*_, providing both mechanistic support and causal interpretability for mediation analysis.

To identify TF_*r*_ → TF_*m*_ → gene_*y*_ pathways, we integrated TF ChIP-seq data from the ENCODE portal [40, 41] with promoter and enhancer annotation to assess binding relationships between TF_*r*_ and TF_*m*_ and between TF_*m*_ and gene_*y*_. The ENCODE K562 collection comprised 749 TF ChIP-seq experiments profiling 524 TFs; only peaks passing the Irreproducible Discovery Rate (IDR) threshold were retained to ensure reproducibility. Promoter regions were defined as -5 kb to +1 kb relative to each gene’s transcription start site, with coordinates obtained from Ensembl [42]. For each TF-gene pair, we assigned a proximal-binding indicator, denoting the presence (1) or absence (0) of a binding relationship based on the overlap between a TF ChIP-seq peak and the gene’s promoter region. Enhancer-gene interaction data were obtained from EnhancerAtlas 2.0 [43], which catalogs 124,879 predicted enhancer-gene links involving 8,800 enhancers and 18,534 genes. For each TF-gene pair, we assigned a distal-binding indicator, denoting the presence (1) or absence (0) of a binding relationship based on the overlap between a TF ChIP-seq peak and an enhancer linked to the gene. By combining proximal and distal binding evidence, we delineated regulatory connections in which the mediator TF_*m*_ is bound by the perturbed TF_*r*_ and binds the outcome gene_*y*_, thereby satisfying the pathway-existence assumption required for causal mediation analysis [24–28].

### Semiparametric model assumptions in CMAPS

To identify the natural direct and indirect effects, we make the following assumptions [34, 45]:

#### Assumption 1

(Causal Consistency) *M* = *M* (*A*) and *Y* = *Y* (*A, M*) = *Y* (*A, M* (*A*)).

Assumption 1 states that the observed values of the mediator and outcome equal their counterfactual values under the observed perturbation status. For example, *M* (1) and *Y* (1,*M* (1)) correspond to the counterfactual expressions of *KLF1* and *HBA1* under *GATA1* perturbation, and are assumed to equal their observed values in *GATA1* -perturbed cells

#### Assumption 2

(Perturbation confounding) For all (*a, a*^*\**^, *m*), *A* ┴ *Y* (*a, m*),*M* (*a*^*\**^) | **C**,*U* and 0 *< p*(*A* = *a* | **C**,*U*) *<* 1.

#### Assumption 3

(Mediator confounding) For all (*a, a*^*\**^, *m*), *M* ┴ *Y* (*a, m*) | *A*, **C**,*U, M* (*a*^*\**^) ┴ *Y* (*a, m*) | **C**,*U*, and 0 *< p*(*m* | **C**,*U*) *<* 1.

The assumptions 2 and 3 state that adjusting for both measured and unmeasured confounders is sufficient to eliminate confounding related to the perturbation indicator and mediator-outcome relationship, respectively. Notably, Assumption 2 is satisfied if sgRNAs are randomly assigned to cells in the perturbation experiment.

Under Assumptions 1–3, we can identify the natural direct and indirect effect as

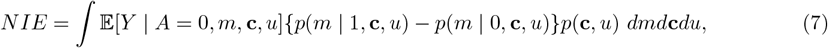

where *p*(·) denotes the probability density or the probability mass function of random variables.

If both *U* and **C** are measured, equation (7) can be applied directly by replacing the expectations and densities with their respective estimates and the integral with its empirical counterpart. However, when *U* is unmeasured, Assumption 4 enables identification of the natural indirect effect (NIE) as *NIE* = *β*_*a*_*θ*_*m*_ [45]. In other words, estimating the NIE requires estimating *θ*_*m*_ and *β*_*a*_ in model (1), which involves both measured confounders **C** and the unmeasured confounder *U*. In contrast, the parameters in model (8) from Baron and Kenny’s method [23] involve only the measured confounders **C**.

Baron-Kenny’s method [23] is the most popular, non-causal mediation analysis, where the goal is to estimate the model parameters ***θ*** and ***β*** from the following equations:

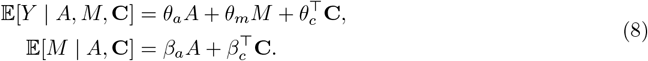

Parameter estimation is typically performed by first regressing the mediator *M* on the exposure *A* and covariates **C** (to estimate *β*), followed by regressing the outcome *Y* on *A, M*, and **C** (to estimate *θ*) [31]. However, a major limitation of this approach is its reliance on the assumption of no unmeasured mediator-outcome confounding (denoted *U*). As noted in previous section, this assumption is often implausible in genomic contexts, where complex regulatory networks introduce numerous sources of unobserved confounding between mediators and outcomes. Moreover, as illustrated in Fig. A4, failure to account for *U* leads to biased estimates of mediation effects and inflated false discovery rates.

We estimate (*θ*_*a*_, *θ*_*m*_, *β*_*a*_) with the semiparametric procedure [34], which evaluates the estimating functions under parametrized models of 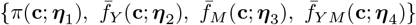, where

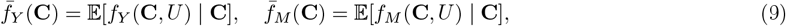

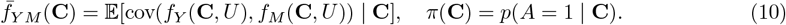

Under regularity conditions, the semiparametric estimator is consistent and asymptotically normal (CAN) [34]. The estimator is multiply robust, provided that at least one of the following model sets is correctly specified: (i) 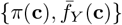, or (ii) 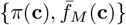, or (iii) 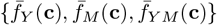. In other words, the estimator remains CAN even when parts of the observed-data model are misspecified. In practice, we model π (**c**) with logistic linear regression and fit 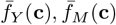, and 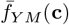with generalized linear models to estimate nuisance parameters (***η***_2_, ***η***_3_, ***η***_4_).

### Adaptive bootstrap test with semiparametric estimator

To obtain valid p-values, we integrate the semiparametric estimator of (*β*_*a*_, *θ*_*m*_) with the adaptive bootstrap (AB) procedure [50]. The AB test statistic is defined as

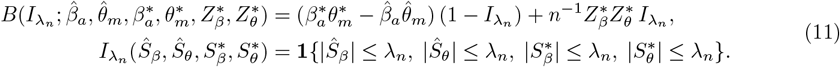

where 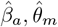 are semiparametric estimates from the observed data, 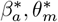are semiparametric estimates from the bootstrapped sample, 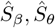and 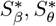are the corresponding standardized statistics, 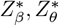 are the corresponding bootstrapped Z-scores, and *λ*_*n*_ is a certain threshold to be specified.

We replace the classical product-of-coefficients estimator of Baron-Kenny’s (BK) method [23] used in the original AB test procedure with the semiparametric estimator because the BK estimator does not account for unmeasured confounding and inflate false discoveries. With this change, both the observed and bootstrapped estimates remain unbiased under the unmeasured mediator-outcome confounding. Theorem 1 shows the non-regular limiting behavior of Walt test statistics at the neighborhood of (*β*_*a*_, *θ*_*m*_) = (0, 0), explaining the conservativeness in our numerical results and in the prior literature [47, 49].

#### Theorem 1

*(i) when* 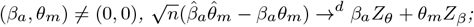

*(ii) when* 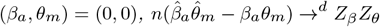

*where* (*Z*_*β*_, *Z*_*θ*_)^*┴*^ ∼ ℬℵ (**0, Σ**), *the asymptotic distribution of semiparametric estimator* 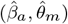.

### Simulation design

Simulations were based on three generative models:

**Setting 1** (A Single Mediator)

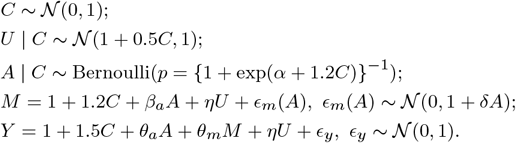

**Setting 2** (Two Mediators)

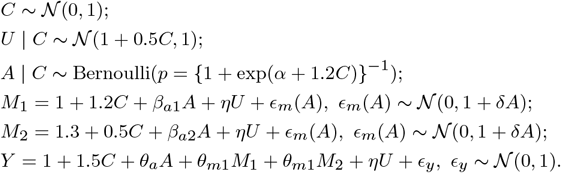

**Setting 3** (Multiple Mediators)

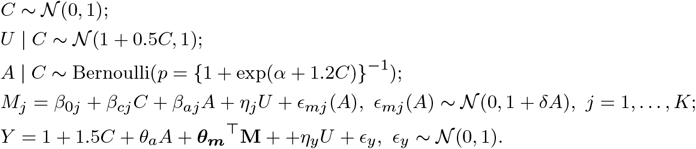

All settings incorporate an unmeasured mediator-outcome confounder *U* and fix *δ* = 5 to induce heteroscedasticity in *M*, thereby ensuring that condition (*) is satisfied.

### Simulation parameters and evaluation metrics

In settings 1 and 2, we vary four factors: (i) sample size, *n* = 5,000 or 10,000; (ii) proportion of treated cells, *p* = 0.05, 0.1, 0.2, or 0.5, controlled via the parameter *α*; (iii) level of unmeasured confounding, *η* = 0 (none) or *η* = 1 (present); and (iv) the values of *β*_*a*_ and *θ*_*m*_, where the mediation effect is *β*_*a*_*θ*_*m*_.Each scenario is replicated 10,000 times.

In setting 3, we fix the sample size at *n* = 5,000 and the proportion of treated cells at *p* = 0.2. For each mediator *j* = 1,&, *K*, we generate *β*_0*j*_∼𝒰 [2, 2] and *β*_*cj*_ ∼𝒰 [1.5, 1.5]. A small subset of the *K* mediators is designated as true mediators with nonzero mediation effects. The remaining mediators have no mediation effect and are evenly assigned to one of three null types: *H*_0,1_: (*β*_*aj*_, *θ* _*mj*_) = (0, 0), *H*_0,2_: (*β*_*aj*_, *θ* _*mj*_) = (1, 0), or *H*_0,3_: (*β*_*aj*_, *θ*_*mj*_) = (0, 1). To induce unmeasured *M*− *Y* dependence, we set *η*_*y*_ = 0.2, with *η*_*j*_ = 0.2 for true mediators and *η*_*j*_ = 0 for null mediators. We vary three factors: (i) the number of mediators, *K* = 30, 50, or 80; (ii) the number of true mediators, 5 or 10; and (iii) the effect size for true mediators, with alternatives *H*_*a*_: (*β*_*aj*_, *θ*_*mj*_) = (0.2, 0.2), (0.2, 0.5), or (1, 1). Each scenario is replicated 5,000 times.

### Perturb-seq in K562 cells

#### Clustering and enrichment analysis

Clustering in Fig. 4a was performed using hierarchical clustering with Euclidean distance and complete linkage, as implemented in the `pheatmap` R package [105]. The input matrix was scaled within each outcome gene across mediators. The optimal number of clusters was determined using the silhouette criterion, which selected six clusters.

Functional enrichment analyses in Fig. 4e for Gene Ontology Biological Process (GOBP), Molecular Function (GOMF), KEGG, and Reactome pathways were conducted using the`clusterProfiler` R package [106].

#### Validation design: negative controls and permutation tests

For the negative control analysis, we used cells transduced with non-targeting (NT) sgRNAs from the EGPS dataset. Among 97 such sgRNAs (cell counts 25-241), we selected three with 190 cells to serve as pseudo-perturbations. For each, we compared one sgRNA against the rest, simulating a two-group design with no expected transcriptomic effect. This design tests CMAPS under a known null condition while maintaining power and group balance. Approximately 8,000 outcome genes were evaluated in each dataset, with each gene tested against an average of 55 mediators.

For the permutation test, we used cells transduced with *GATA1* perturbation and NT sgRNAs and independently permuted treatment assignments *A*, mediators *M*, and outcomes *Y*, thereby disrupting the causal pathway and ensuring *NIE* = *β*_*a*_*θ*_*m*_ = 0. This synthetic null was evaluated on the same scale with approximately 8,000 outcome genes per replicate, each tested against an average of 69 mediators.

#### Validation using the inspre causal regulatory network

We applied inspre [21] to infer causal regulatory relationships among *GATA1*, the outcome gene *BCL2L1*, and candidate mediators supported by regulatory evidence. Within a regression-based framework, inspre models perturbation effects by jointly analyzing gene expression of all genes in the network measured across cells subjected to each gene-targeting perturbation in the set, together with control cells.

For the (*GATA1, BCL2L1*) pair, 86 candidate mediators were defined based on regulatory evidence. We restricted the analysis to mediators with valid perturbations in the K562 Perturb-seq data, defined as targeting sgRNAs that achieved at least a 30% reduction in target gene expression relative to control cells and were represented by at least 50 perturbed cells. Inspre was therefore applied to *GATA1, BCL2L1*, and the 12 mediators that satisfied these criteria, including *STAT5A*, which was identified as statistically significant by CMAPS.

Inspre infers regulatory networks using *L*_1_-regularized regression to induce sparsity and select a minimal set of regulatory relationships. Increasing the *L*_1_ penalty yields sparser networks by shrinking regression coefficients toward zero. After model fitting, edges are further pruned based on the absolute magnitude of the estimated coefficients to remove weak or unstable regulatory relationships, with the sign of each retained coefficient indicating the inferred direction of regulation (positive or negative). Network sparsity was defined as the proportion of non-zero edges among all possible edges. As shown in Fig. A7, varying the *L*_1_ penalty and the minimum edge-strength threshold produced nine networks with sparsity ranging from 0.03 to 0.22, enabling assessment of sensitivity to parameter choices. The network shown in Fig. 4b corresponds to an intermediate sparsity level (0.14) within this range.

### MultiPerturb-seq in AT/RT-derived BT16 cells

#### Data preprocessing

We analyzed paired scRNA-seq and scATAC-seq profiles generated from MultiPerturb-seq [35], a pooled CRISPR interference screen conducted in BT16 cells. Following the original study’s quality-control cfriteria, we retained cells with at least 100 detected RNA transcripts or 100 ATAC-seq fragments. The screen targeted 99 epigenetic remodelers, with per-target cell counts ranging from 1 to 7,895 (median: 578). An additional 4,528 cells carried non-targeting sgRNAs and served as controls. To ensure sufficient power and balanced treatment-control groups, we restricted causal mediation analysis to the 73 perturbations represented by at least 190 cells (Fig. A6a). Each of these perturbations was analyzed alongside the pool of non-targeting control cells, resulting in datasets where treated cells comprised at least 4% of the total sample.

The scRNA-seq data included gene-level transcript counts for 25,217 genes, and the scATAC-seq data contained fragment counts across 155,949 peaks. In the original study, each ATAC-seq peak was annotated to one of 11 genomic features and mapped to a gene. In total, 25,077 genes were linked to at least one ATAC peak, with 95% of genes associated with 15 or fewer peaks. To prepare inputs for CMAPS analysis, we retained genes with matched measurements in both RNA and ATAC modalities. Genes were harmonized using HGNC symbols and Ensembl gene identifiers, resulting in 18,226 genes with both gene expression and chromatin accessibility data.

For chromatin accessibility mediators, we adopted the 11-category peak annotations reported from the original study and grouped them into four broad *cis*-regulatory element classes: promoter, proximal, distal intergenic, and intragenic (Table 1). For each gene in each cell, CRE-level accessibility was calculated by summing the raw fragment counts of all peaks mapped to that gene within the same CRE group.

Both the gene-expression and CRE-accessibility matrices were normalized using the *LogNormalize* method from the `Seurat` R packages [107]. Briefly, raw counts were divided by the total counts per cell, multiplied by a scale factor of 10^6^, and then natural-log transformed. The resulting normalized matrices served as input for downstream CMAPS analysis.

#### CMAPS analysis and gene filtering

For each perturbation, CMAPS was applied to model the effect of perturbation on gene expression mediated through CRE accessibility. Each analysis included treated cells with sgRNAs targeting a specific gene and control cells carrying non-targeting sgRNAs. Genes were excluded from a given perturbation if their expression was constant across cells. Similarly, CRE-gene pairs were excluded if the gene’s CRE accessibility was constant. The number of genes and CRE-gene pairs analyzed varied across perturbations, with an average of 13,231 genes and 20,275 CRE-gene pairs per perturbation. Among the analyzed genes, 83% had promoter peaks, 38% proximal, 24% distal intergenic, and 7% intragenic peaks. The distribution of analyzed genes and CRE-gene pairs per perturbation is shown in Fig. A6b-c.

#### Clustering and enrichment analysis

Clustering in Fig. 5a was performed using hierarchical clustering with Euclidean distance and complete linkage, as implemented in the `pheatmap` R package [105]. The input matrix was not scaled, as the values represent proportions that sum to one across each row. The optimal number of clusters was determined using the silhouette criterion, which selected three clusters.

Functional enrichment analyses in Fig. 5d for Gene Ontology Biological Process (GOBP), Molecular Function (GOMF), KEGG, and Reactome pathways were conducted using the `clusterProfiler` R package [106]. We did not apply false discovery rate (FDR) correction because the gene universe used for enrichment comprised only 73 perturbation target genes, making enrichment tests highly discrete and limiting the number of meaningful hypotheses. In such cases, multiple testing correction methods can be overly conservative, leading to the omission of biologically relevant signals. Instead, we prioritized interpretability and hypothesis generation by selecting enriched terms with nominal p-values *<* 0.1 and fold enrichment ≥1.2. We acknowledge that this approach increases the potential for false positives and interpret these findings accordingly.

## Notes

### Competing Interest Statement

The authors have declared no competing interest.

