## Supplemental Table A1 and Figures A1-7 for "CMAPS: Causal Mediation Analysis of Perturbation Screens with Application to Genome-scale Perturb-seq Data"

### Appendix A Supplementary Materials

| Method | Target task |
|---|---|
| MIMOSCA [10] | Effects of multiple perturbations on gene expression |
| scMAGeCK [11] | Differential expression analysis combined with Robust Rank Aggregation |
| SCEPTRE [12] | Differential expression analysis combined with conditional randomization test |
| Normalisr [13] | Differential expression analysis with integrated normalization and association testing |
| GPSA [14] | Identification of candidate causal perturbations from differential gene expression data |
| GSFA [15] | Differential expression analysis combined with Bayesian factor analysis |
| Mixscale [16] | Differential expression analysis with modeling of perturbation efficiency |
| MUSIC [17] | Model-based prioritization of gene perturbation effects |
| Energy test [9] | Test for distributional differences between perturbed and control cells |
| CINEMA-OT [18] | Perturbation effects via optimal transport matching of counterfactual cell pairs |
| STE [19] | Differential expression analysis via standardized average and quantile treatment effects |
| RCSP [108] | Identify causal ordering of gene expression |
| RENGE [20] | Infer gene regulatory networks |
| inspre [21] | Infer causal regulatory networks |
| CausalBench [22] | Benchmark for gene regulatory networks |

**Table A1** Summary of existing methods for the analysis of perturbation experiments and their associated target tasks.

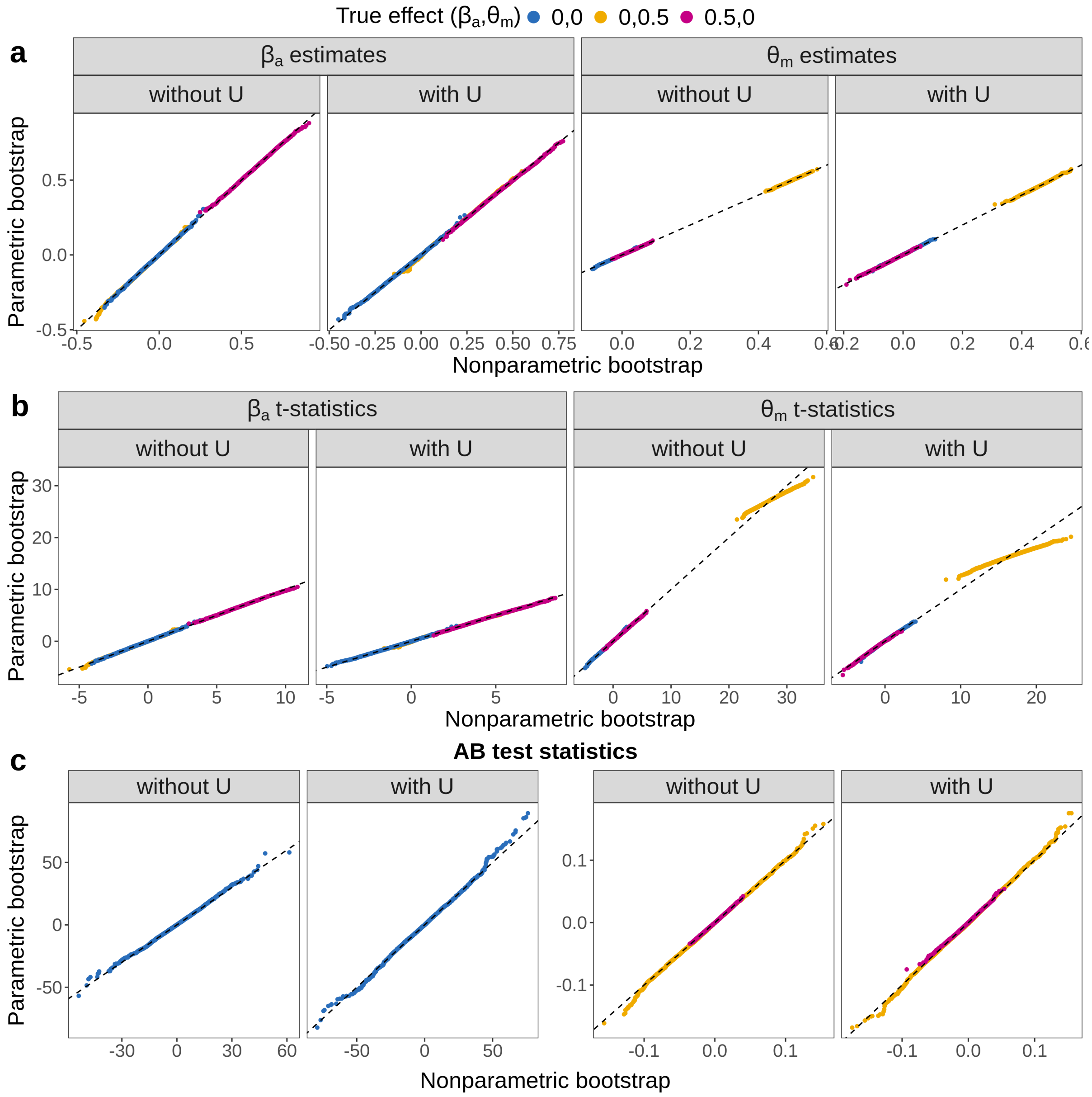


**Fig. A1** Comparison of 10,000 sorted nonparametric bootstrap replicates with their corresponding parametric bootstrap replicates under three null hypothesis settings, both without and with the unmeasured mediator-outcome confounding $U$. **a** Empirical distribution of estimates. **b** Empirical distribution of standardized statistics. **c** Empirical distribution of adaptive bootstrap (AB) test statistics.

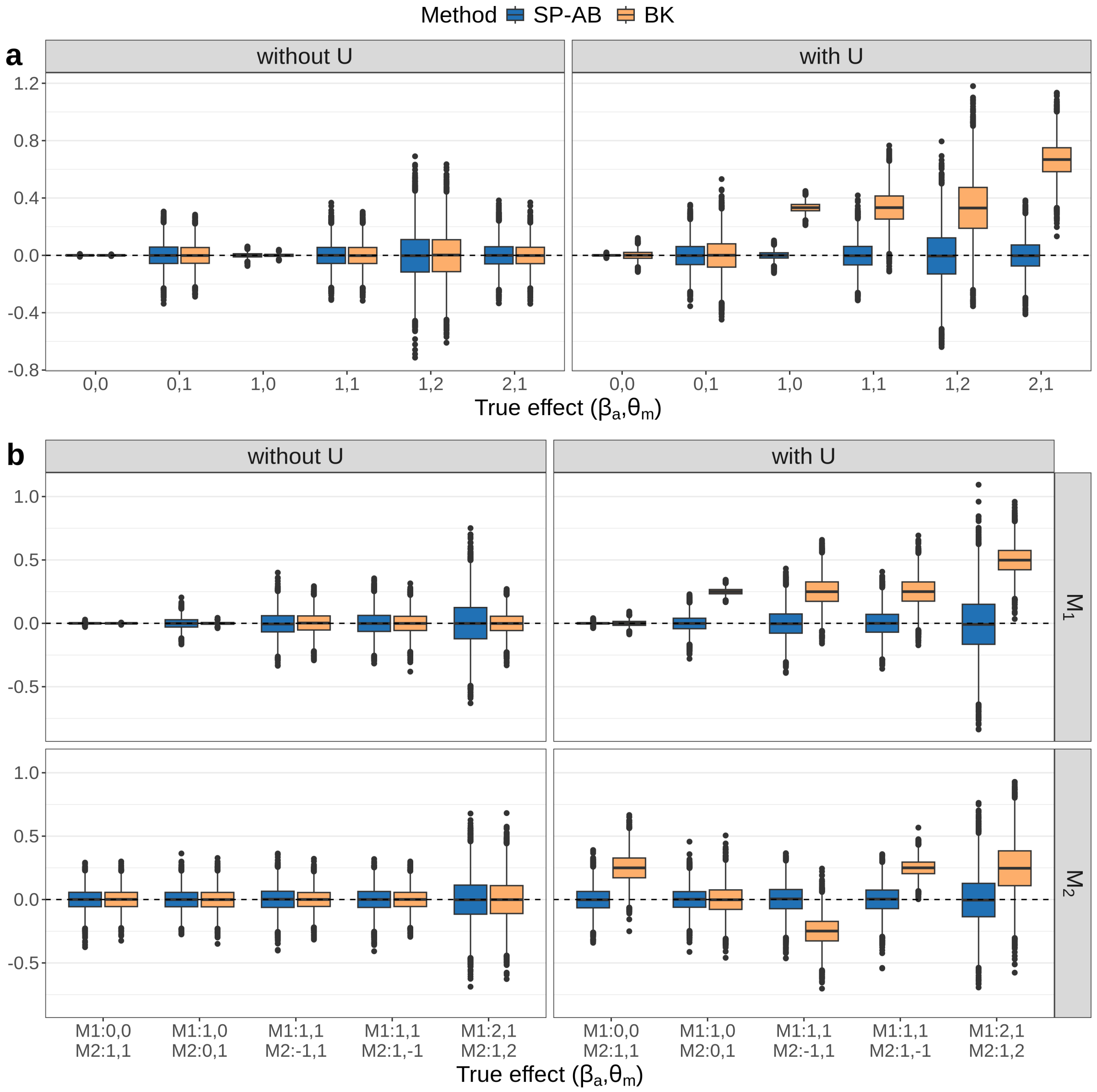


**Fig. A2** Estimation accuracy of NIE estimates under varying true mediation effects. **a** Boxplots of estimation error (difference between estimated and true effects) in the single mediator setting (Setting 1). **b** Boxplots of estimation error in the two-mediator setting (Setting 2); each row corresponds to a distinct mediator.

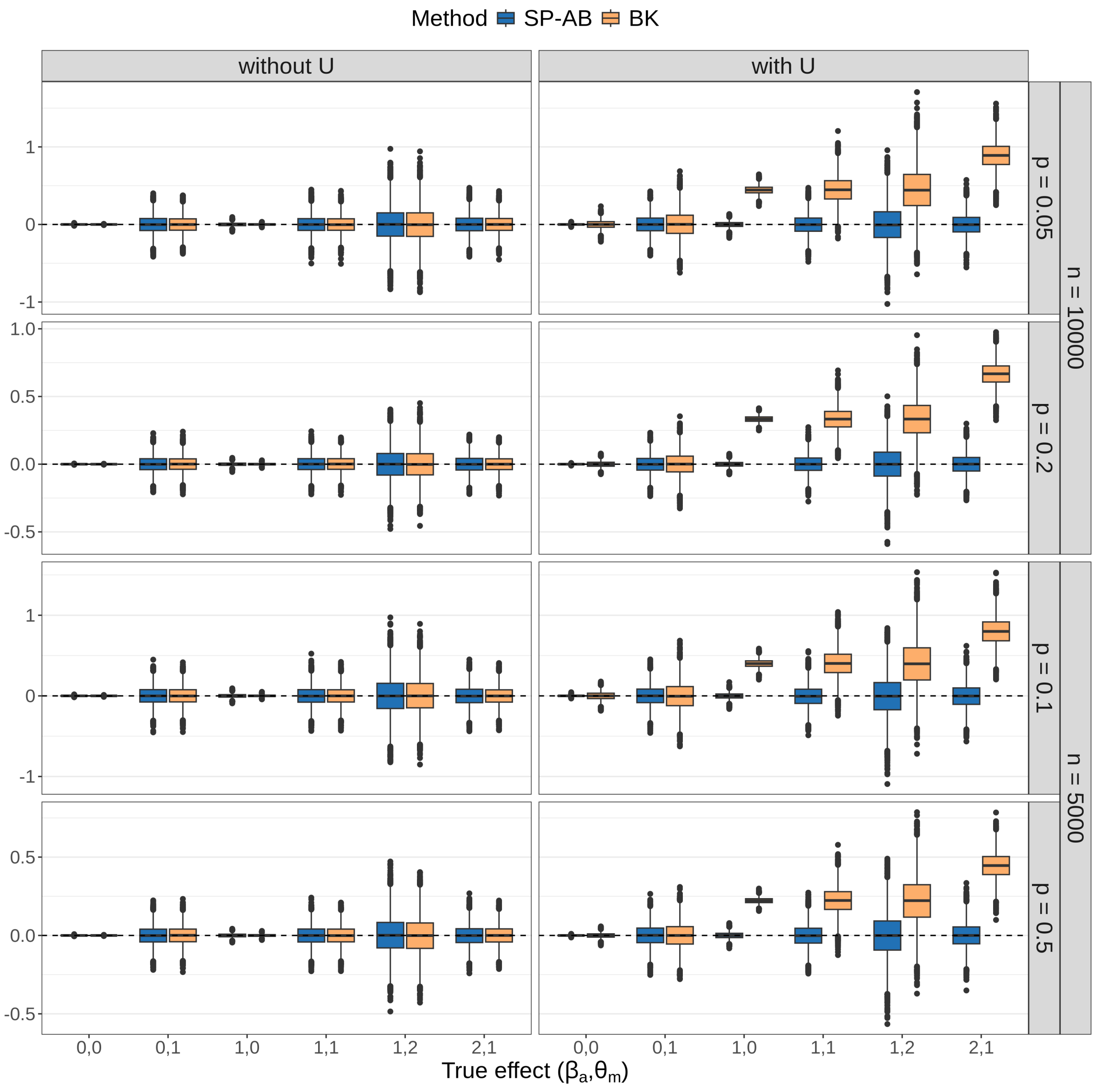


**Fig. A3** Boxplots of estimation error of NIE (estimated minus true NIE) under varying sample sizes and treatment proportions in the single-mediator setting. Columns indicate the absence or presence of unmeasured mediator-outcome confounding. Rows correspond to combinations of sample size and proportion of treated cells.

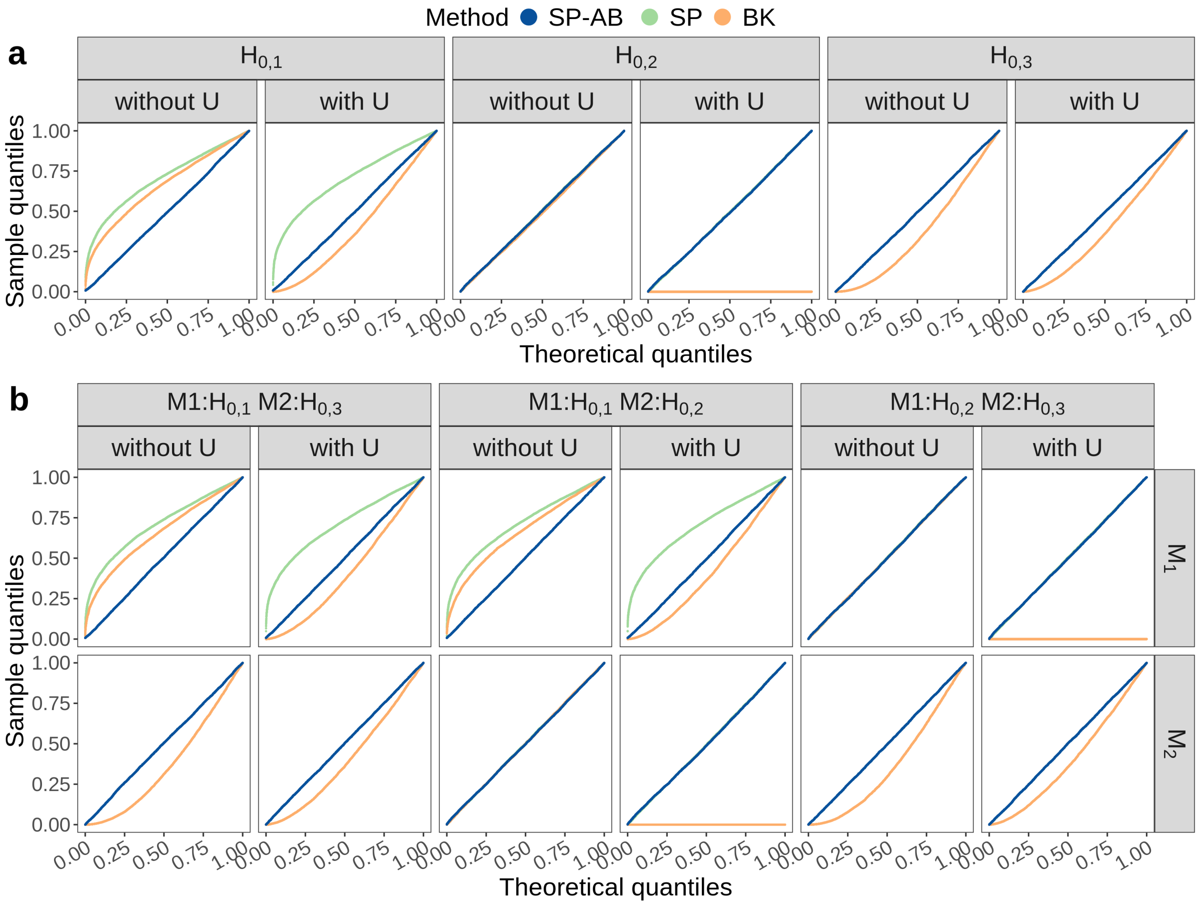


**Fig. A4** Q-Q plots comparing the empirical p-value distributions with the theoretical Uniform(0,1) null distribution under three types of null hypotheses: $H_{0,1} : (\beta_a, \theta_m) = (0, 0)$, $H_{0,2} : (\beta_a, \theta_m) = (1, 0)$, and $H_{0,3} : (\beta_a, \theta_m) = (0, 1)$. **a** Q-Q plots of p-values in the single mediator setting (Setting 1). **b** Q-Q plots of p-values in the two-mediator setting (Setting 2), with rows representing the two mediators, each tested under a different null type.

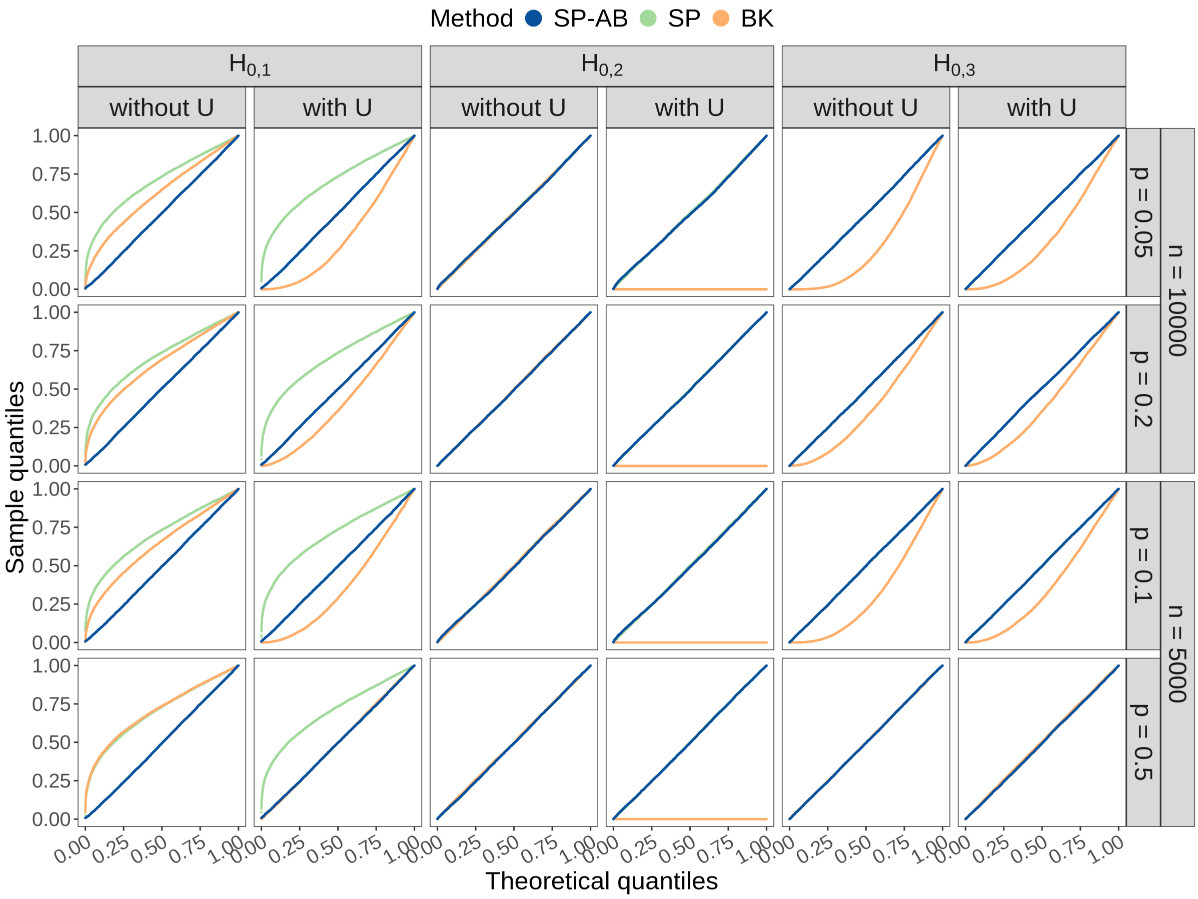


**Fig. A5** Q-Q plots of p-values under varying sample sizes and treatment proportions in the single-mediator setting. Theoretical quantiles are those of the Uniform(0,1) null distribution. Columns indicate three types of null hypotheses: $H_{0,1} : (\beta_a, \theta_m) = (0, 0)$, $H_{0,2} : (\beta_a, \theta_m) = (1, 0)$, and $H_{0,3} : (\beta_a, \theta_m) = (0, 1)$. Rows correspond to combinations of sample size and proportion of treated cells.

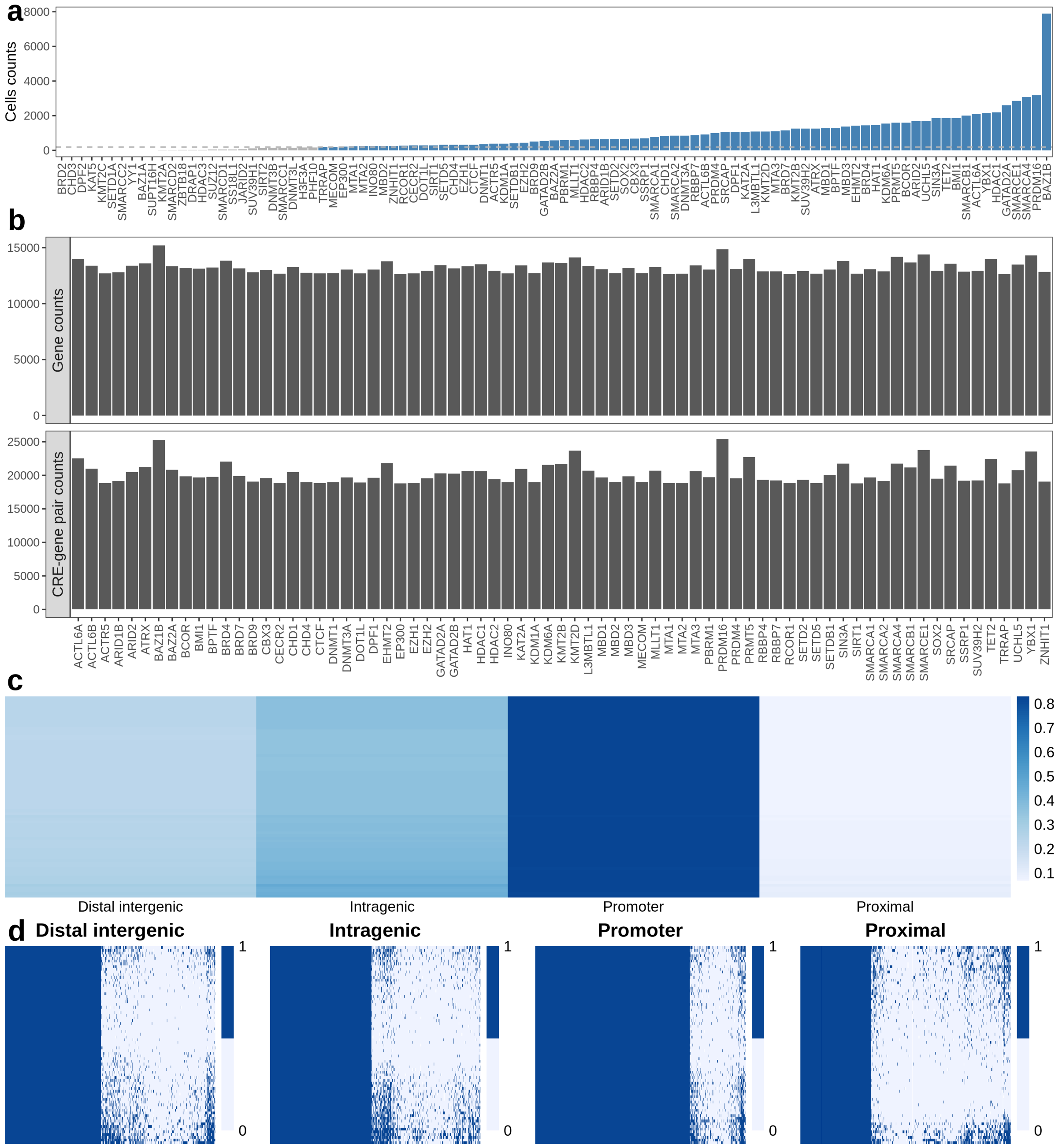


**Fig. A6** Overview of perturbation-level data filtering and inclusion criteria for CMAPS analysis. **a** Number of cells per perturbation across 99 targeted epigenetic remodelers. Each bar represents a gene target. The horizontal dashed line denotes the inclusion threshold of 190 cells. 73 perturbations (blue) met this criterion and were included in downstream CMAPS analysis, whereas excluded perturbations are shown in gray. **b** Number of genes (top) and CRE-gene pairs (bottom) analyzed per perturbation after filtering for nonzero variance in expression and accessibility, respectively, across the 73 included perturbations. **c** Heatmap showing the proportion of analyzed genes carrying at least one peak in each CRE category per perturbation. Rows represent the 73 included perturbations; columns correspond to the four CRE groups. **d** Binary heatmaps of gene-CRE peak presence for each of the four CRE categories. Rows represent the 73 included perturbations; columns represent genes. A value of 1 indicates the presence of at least one peak for the gene in the corresponding CRE under the perturbation. We observed no discernible patterns in the binary presence of peaks across genes and perturbations, indicating that the variation in CRE peak proportions in Fig. 5a reflects genuine biological differences. This supports the validity of our downstream enrichment analysis.

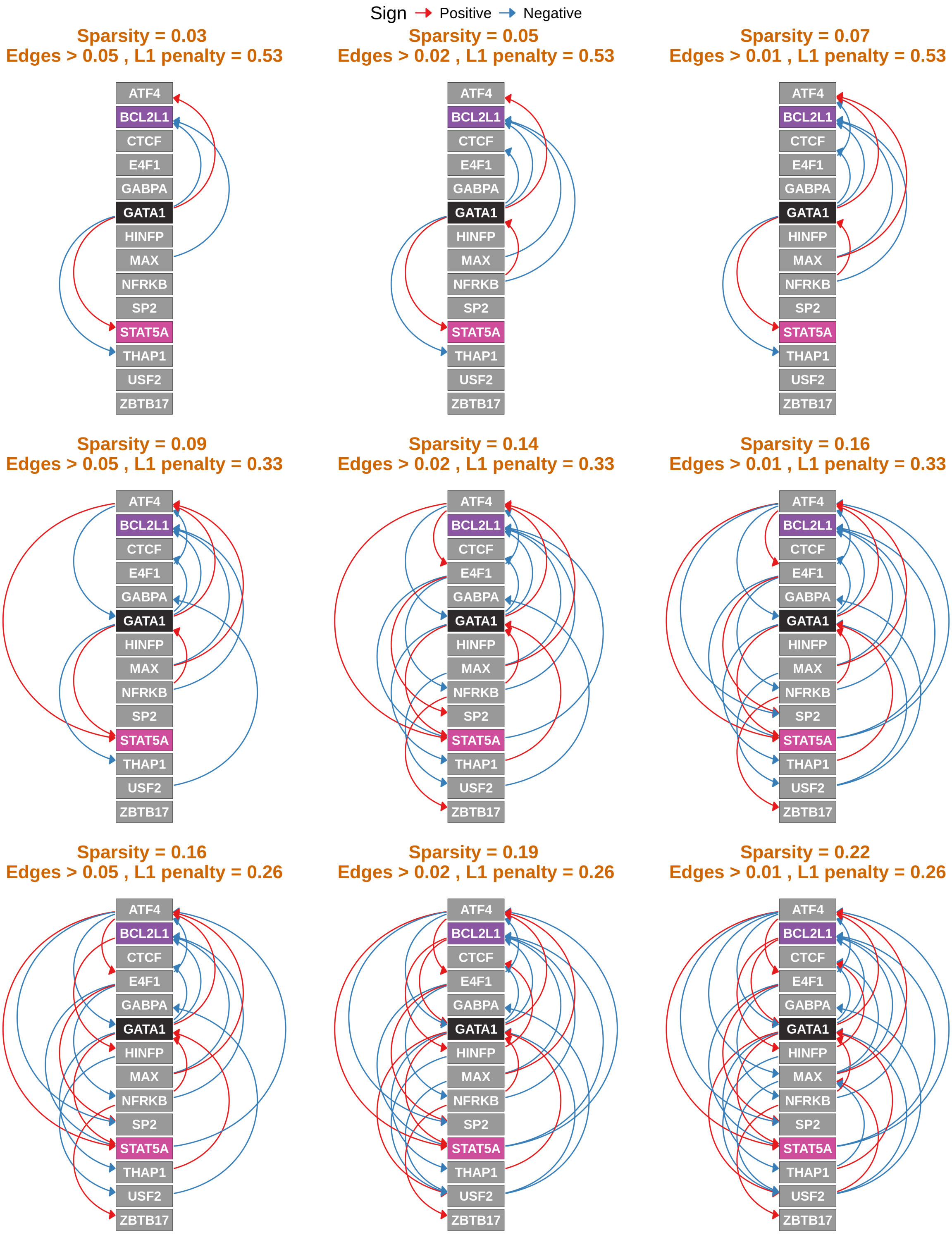


**Fig. A7** Inspre-inferred regulatory networks under varying pruning parameters. Nine networks inferred by inspre using different $L_1$ penalties and minimum edge-strength thresholds are shown; *GATA1*, *BCL2L1*, and *STAT5A* are highlighted, and red and blue edges indicate positive and negative regulation, respectively.
